# Adaptive evolutionary trajectories in complexity: repeated transitions between unicellularity and differentiated multicellularity

**DOI:** 10.1101/2024.05.14.594091

**Authors:** Hanna Isaksson, Peter Lind, Eric Libby

## Abstract

Multicellularity spans a wide gamut in terms of complexity, from simple clonal clusters of cells to large-scale organisms composed of differentiated cells and tissues. While recent experiments have demonstrated that simple forms of multicellularity can readily evolve in response to different selective pressures, it is unknown if continued exposure to those same selective pressures will result in the evolution of increased multicellular complexity. We use mathematical models to consider the adaptive trajectories of unicellular organisms exposed to periodic bouts of abiotic stress, such as drought or antibiotics. Populations can improve survival in response to the stress by evolving multicellularity or cell differentiation—or both; however, these responses have associated costs when the stress is absent. We define a parameter space of fitness-relevant traits and identify where multicellularity, differentiation, or their combination is fittest. We then study the effects of adaptation by allowing populations to fix mutations that improve their fitness. We find that while the same mutation can be beneficial to phenotypes with different complexity, e.g. unicellularity and differentiated multicellularity, the magnitudes of their effects can differ and alter which phenotype is fittest. As a result, we observe adaptive trajectories that gain and lose complexity. We also show that the order of mutations, historical contingency, can cause some transitions to be permanent in the absence of neutral evolution. Ultimately, we find that continued exposure to a selective driver for multicellularity can either lead to increasing complexity or a return to unicellularity.

## Introduction

There is a broad spectrum of multicellularity that ranges from primitive replicating groups of cells to large complex organisms with intricate tissues and behaviors (1–5). Recent experimental studies have demonstrated that many different unicellular organisms can readily evolve simple forms of multicellularity, in response to a variety of selective pressures (6–10). However, these forms of multicellularity are often only one mutation away from unicellularity (11) and lack the basic features of fully fledged multicellular organisms such as cell adhesion and cell-cell communication (12–14). While it may simply be a matter of time before the same selection for multicellularity eventually drives increased complexity, such as cell differentiation, other possibilities exist as well. For example, selection may favor multicellularity and/or cell differentiation but not both together, e.g. choanoflagellates express multiple cell types across their life cycle but when forming multicellular colonies all cells are of the same type (15, 16). Further complicating matters is the possibility that adaptation may render the benefits conferred by multicellularity or cell differentiation superfluous, e.g. adaptation to a toxin may lead to resistance which nullifies any gains in survival offered by forming multicellular groups. Here, we consider the question of whether a single type of selection is sufficient to sustain the evolution from unicellularity to differentiated multicellularity.

Previous studies of the evolution of differentiated multicellularity have focused on identifying which innovation came first: differentiation or multicellularity. In support of differentiation first, many unicellular organisms have the ability to express different phenotypes, or cell types, in order to switch between tasks in time (17, 18). During the subsequent evolution of multicellularity the different cell types could then be arranged in spatial patterns, co-opted for new functions by the nascent multicellular organism (19–21). Another possibility is that multicellularity evolved first and then generated a new context that allowed the evolution of de novo cell types. For some model systems this involved modification of a trait already present in unicellular populations, e.g. increasing cell death rate (22, 23); but in other model systems differentiation evolved by partitioning two tasks typically performed simultaneously in unicellular organisms across two different cells (24, 25). Establishing the order of innovations can be useful in both understanding the early drivers of differentiated multicellularity and distinguishing between the resulting forms, but we currently lack models that allow us to determine when each path is more likely to occur.

Even if there is a path from unicellularity to differentiated multicellularity it is unclear to what extent it can be maintained. During the transition to multicellularity cells evolved higher levels of cooperation (26, 27). A common issue faced during the evolution cooperation is the threat that cheating mutants may arise and ultimately undermine the cooperation (28–30). The costly effects of cheating can be mitigated by certain population structures, including the organization of cells into multicellular groups (31). Theoretical models have explored how cooperation, in the form of differentiation, may evolve in simple multicellular organisms (32, 33). These models often assume that populations are multicellular and then assess the value of differentiation within a multicellular context (21, 34, 35). Yet, early forms of multicellularity are often only a mutation away from unicellularity (11), and so reversion back to unicellularity is a constant threat (36– 38). Moreover, adaptations could reduce the benefits conferred by differentiation and/or multicellularity and also lead to reversions. For example, in response to antibiotics bacterial populations can improve their survival by differentiating into special survival phenotypes or by forming multicellular biofilms (39, 40). Should the bacteria evolve resistance to the antibiotic it would reduce the benefits of both differentiation and multicellularity and lead to a loss in complexity.

The example of antibiotics as a driver for both the evolution of complexity and its loss may be more generally applicable. We can more broadly categorize antibiotics as a type of abiotic stress where “abiotic” signifies that the stress does not evolve in response to its effects. Other examples of abiotic stress include desiccation (41–43), high and low pH (44–46), salt (16, 47), etc, all of which threaten the survival of unicellular organisms but do not co-evolve with them. We draw a distinction with biotic stress, e.g. predation, in which co-evolution of two species may lead to changes in responses over time, e.g. the well known arms races (48, 49). Abiotic stress has been identified as a potent selector for the evolution of multicellularity (50). In addition to multicellularity, abiotic stress may also select for types of differentiation. Bacterial persistence and bet-hedging more generally have been shown to be favored in environments with regular bouts of abiotic stress (51–56).

Here we study the evolutionary trajectory from unicellularity towards differentiated multicellularity when a population is repeatedly exposed to an abiotic stress. In response to the stress cells may improve survival by forming multicellular groups or differentiating into survival specialists, or both. We use a mathematical modeling framework to weigh the benefits of survival from these complex phenotypes versus the costs they impose to growth, manifesting as time delays as cells switch between types or forms. We also explore the stability of the evolved phenotypes and whether further adaptation leads to increasing complexity or reversion to unicellularity. Through our analyses we find that both multicellularity and cell differentiation can readily evolve under the same abiotic stress, but not necessarily in the same life cycle. We also find that the path taken and its stability depend heavily on historical contingency and stochasticity, with the possibility of adaptive gains and losses of complexity. Thus we conclude that abiotic stress may well drive the evolution of differentiated multicellularity, but it may only appear in transient episodes along a path that ultimately returns to unicellularity.

## Results

Our aim is to explore the evolutionary paths leading from unicellularity to differentiated multicellularity when the selective driver is a type of abiotic stress. In particular we are interested in what route towards complex multicellularity is the most likely i.e. whether differentiation or group formation evolves first, and how adaptation affects the trajectory and stability of evolution, see Fig. 1A. We consider a model in which a population of cells experiences an environment that fluctuates between two states: 1. a neutral state (*E*_0_) where cell populations grow exponentially according to a rate *b* and 2. a state containing an abiotic stress (*E*_*A*_) that prevents populations from growing and causes them to die at a rate *d*. In this model the fitness of a population depends on the product of growth in *E*_0_ and survival in *E*_*A*_. For simplicity, we assume that the time spent in each environment is the same, similar to the approach in (57). In the subsequent sections we consider what conditions favor the evolution of cell differentiation, multicellularity, or both.

**Fig. 1.**
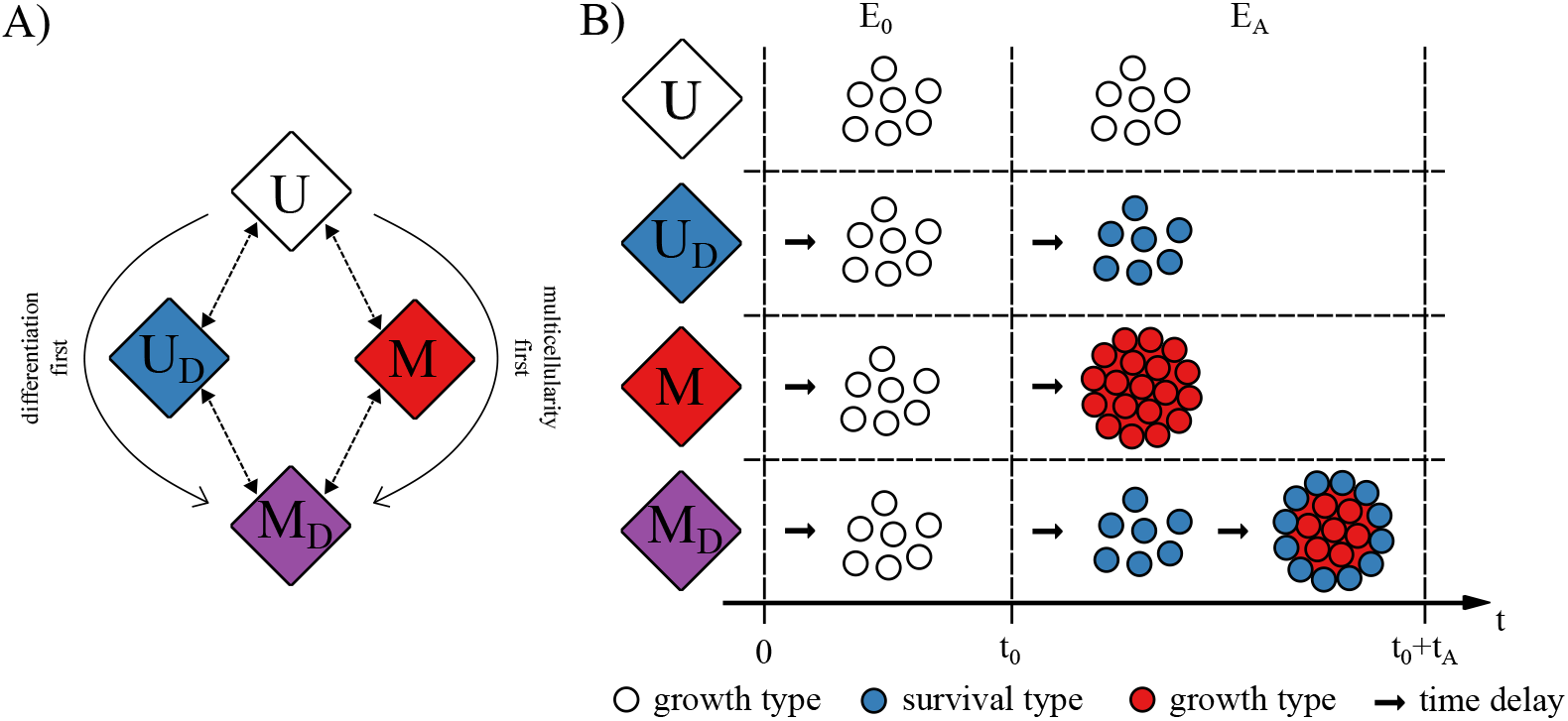
Evolutionary paths to differentiated multicellularity. A) A schematic shows two different evolutionary routes from unicellularity (*U* ) to differentiated multicellularity (*M*_*D*_) depending on whether unicellular populations evolve differentiation or multicellularity first. B) A schematic shows how different populations respond to fluctuations between environments without stress (*E*_0_ ) and environments with an abiotic stress (*E*_*A*_). The strictly unicellular population (*U*, white cells) grows during *E*_0_ but then dies during exposure to the stress *E*_*A*_. By differentiating into a survival phenotype (*U*_*D*_, blue cells) cells become resistant to the stress. Another way to increase survival in response to the stress is to form multicellular groups (*M*, red cells), where the outer cells provide physical protection for inner cells. Differentiation and group formation can also be combined in a joint lifestyle, where cells form groups and the outer cells differentiate (*M*_*D*_, blue and red cells). Each stress response comes with a cost in the form of time delays (represented by arrows).

### Differentiated unicellularity

Cells can improve their fitness by evolving strategies that better protect them from the stress, see Fig. 1B. One way for cells to do this is through differentiation into a specialized survival phenotype in the stressful environment. We assume that the survival phenotype allows cells to completely survive the stress but in this differentiated state they cannot reproduce — analogous to the flagellation constraint in Volvocales where cell division and flagellated motility can not take place simultaneously (58). We also assume that differentiating between phenotypes comes with a cost to cell growth. The cost manifests as a time lag *τ*_*D*_ when cells switch phenotypes; so when the environment switches states from stressful to neutral, there is a mismatch between the phenotype and the environment leading to a temporary reduction in growth (59, 60), see Fig. 2A. Thus, the fitness of differentiated unicellularity is determined by the interplay between two factors: the birth-death ratio and the time delay from switching phenotypes.

**Fig. 2.**
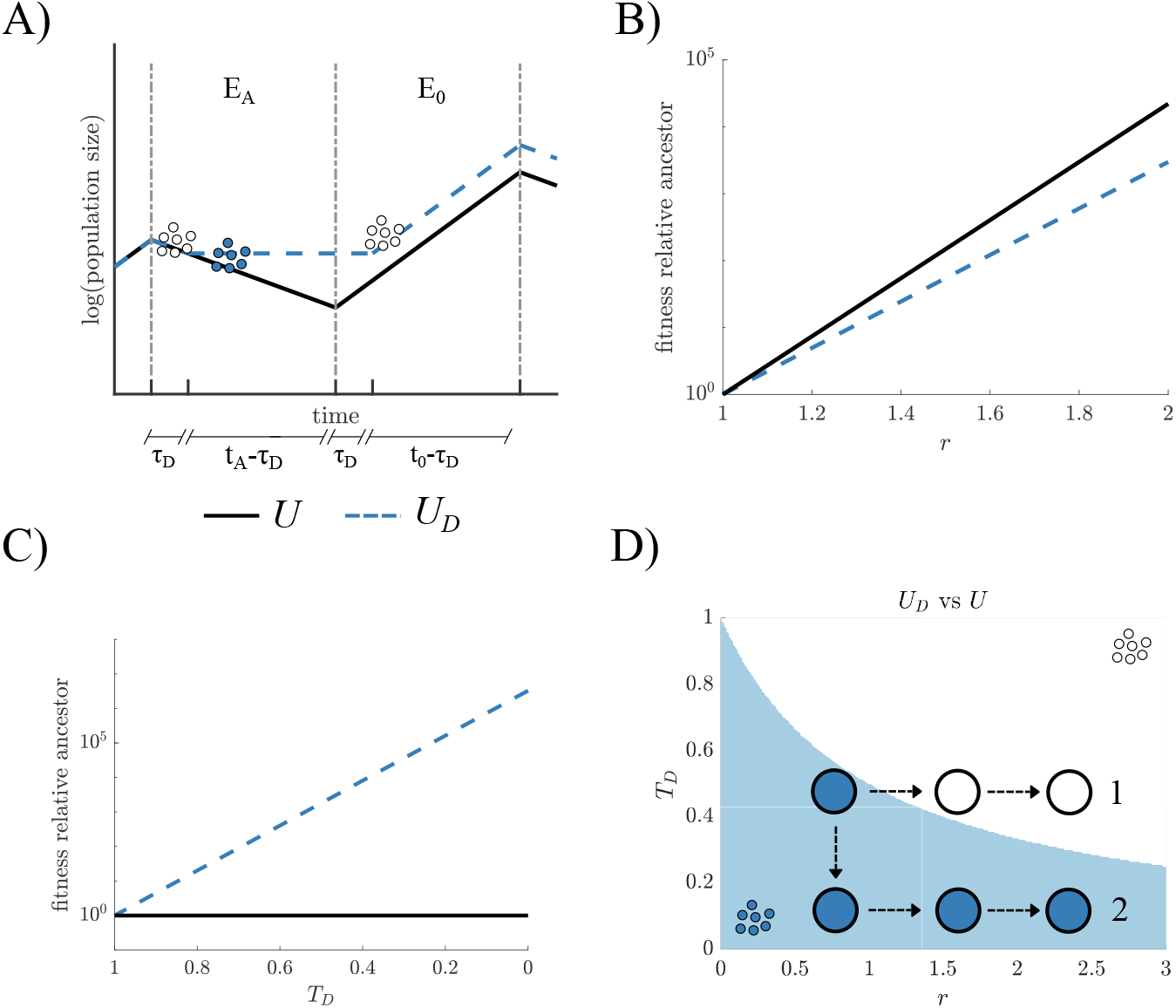
Fitness advantage of differentiation and evolutionary trajectories. A) A schematic illustrates the dynamics of two populations in an environment that switches between two states: with and without an abiotic stress, *E*_*A*_ and *E*_0_ respectively. The first population is unicellular (*U*, solid line) which grows exponentially in *E*_0_ and decays in *E*_*A*_. The second population (*U*_*D*_, blue dashed line) is similar to the first except that it differentiates to a stress-resistant phenotype that survives the stress in *E*_*A*_ but can not reproduce. Differentiated populations experience a time lag *τ*_*D*_ each time they switch to and from the stress-resistant phenotype. B) The two lines show how beneficial mutations in the birth-death ratio (*r*) change the fitness relative to an ancestor (*r* = 1) in *U* (solid line) and *U*_*D*_ (blue dashed line) populations. Beneficial mutations in *r* always cause larger fitness gain in *U* than *U*_*D*_. C) A plot shows the effects of beneficial mutations in the normalized time delay *T*_*D*_ = *τ*_*D*_*/t*_0_. Decreasing *T*_*D*_ increases the fitness of *U*_*D*_ relative to its ancestor (*T*_*D*_ = 1), but has no effect on *U* . D) A parameter space shows the values of the traits *r* and *T*_*D*_ and is colored according to which is fitter *U* (white) or *U*_*D*_ (blue). Populations undergo adaptive walks in parameter space via beneficial mutations in *T*_*D*_ and *r*. Two example trajectories show that the type and order of mutations determine whether a *U*_*D*_ population will maintain differentiation (trajectory 2) or cross into a region of the parameter space where loss of differentiation is favored and mutations in *T*_*D*_ are neutral (trajectory 1).

Using a simple modeling framework of exponentially growing populations (see Methods: Equations for population growth), we can identify the conditions for which differentiation confers a fitness advantage, i.e. differentiated unicellularity (denoted *U*_*D*_) is fitter than undifferentiated unicellularity (denoted *U* ). We let *r* represent the birth-death ratio and *T*_*D*_ represent the fraction of time in *E*_0_ when cells do not grow because they are still in the survival state — we note that *T*_*D*_ is *τ*_*D*_ scaled by the time spent in *E*_0_. We then find that differentiation provides a fitness advantage when

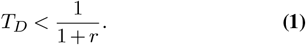

Eq. (1) shows that the value of *r* limits which time lags allow differentiation to be fitter. Since *r* is a ratio of the population growth rate in *E*_0_ and the death rate in *E*_*A*_, it encapsulates both the loss in growth and the gain in survival from switching. If *r* is low, either because populations grow slowly without stress or because the abiotic stress is particularly challenging, differentiation with high time lags can be advantageous.

One consequence of Eq. (1) is that adaptation can alter whether differentiation is favored. If we consider adaptation in terms of mutations that increase the value of *r*, either by improving the growth rate in *E*_0_ or survival in response to the abiotic stress, fitness increases in both *U* and *U*_*D*_, see Fig. 2B. However, in relative terms, *r* mutations increase fitness more in *U* so that continually increasing *r* will eventually cause unicellularity to become fitter than differentiation. In contrast, if we consider adaptation in terms of mutations that lower the time delay in switching this only has a fitness effect on *U*_*D*_ life cycles, see Fig. 2C. Lowering the time delay allows differentiation to be favored even for large values of *r*.

A result of these findings is that although differentiated unicellularity can improve fitness by either mutations in *r* or *T*_*D*_, the order of these mutations can alter whether differentiation is maintained or lost. Figure 2D shows that a mutation that lowers the time delay (resulting in lower *T*_*D*_) followed by one that increases the birth/death ratio *r* can keep an organism with differentiation in areas of parameter space (*r, T*_*D*_) where differentiation is fittest. Yet, if a mutation occurs in *r* first it can lead the population into an area of parameter space where unicellularity is fittest. A reversion to unicellularity would then mean that mutations in *T*_*D*_ are neutral and may not fix, preventing the population from evolving to an area of parameter space where differentiation is fitter.

### Multicellularity

Another way for cells to improve their fitness during stress (*E*_*A*_) is by forming multicellular groups, which provide physical protection to some interior set of cells (50, 61, 62). We choose to focus on multicellularity formed via aggregation, since this group forming strategy responds faster to environmental changes than clonal multicellularity (63). We assume that in the aggregated group, cells are partitioned into exterior and interior compartments such that cells on the exterior directly experience the abiotic stress while those in the interior are temporarily shielded from the stress. In the stressful environment neither external nor internal cells can reproduce, which is common in aggregative multicellularity (64). For simplicity, we also assume that upon exposure to the stress exterior cells die at the same rate as if they were unicellular and then are immediately replaced by interior cells until there are no interior cells left. Thus, forming multicellular groups does not offer permanent protection but it does reduce the population death rate. We characterize different types of groups by introducing a parameter *R* which is the initial ratio of the number of interior cells to exterior cells; higher values of *R* indicate structures with a higher fraction of protected interior cells. Since we do not explicitly impose a type of topology the physical structure of the group can vary across *R* values, see Fig. 3A. Finally, we assume that multicellularity has a cost, similar to differentiation, which appears as a time lag *τ*_*G*_ between forming and breaking apart groups; so as in the case of differentiation there is a trade-off between increased survival in *E*_*A*_ and missed growth opportunities in *E*_0_ (for details see Methods: Equations for population growth).

**Fig. 3.**
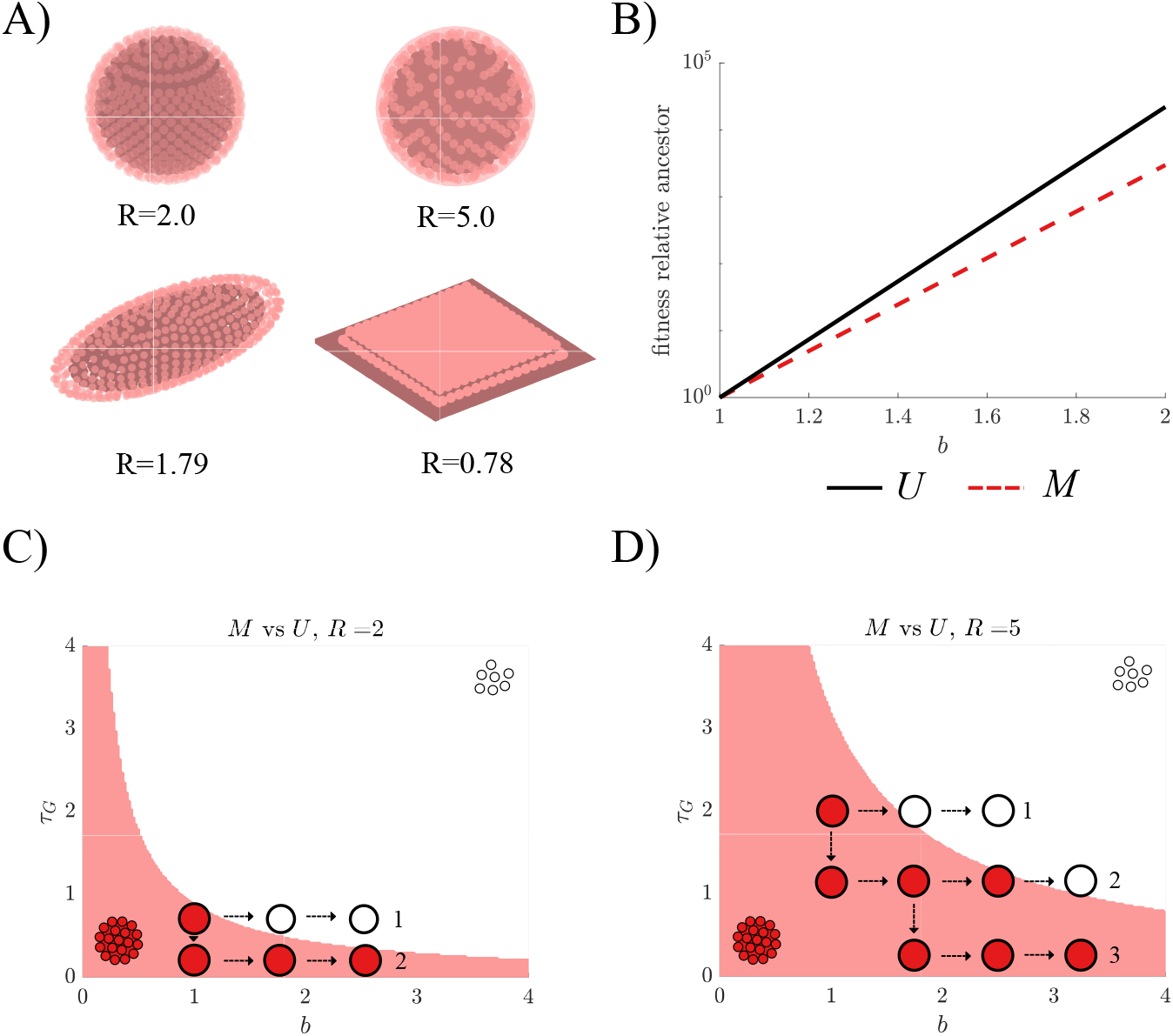
Evolution of multicellularity depends on group topology and order of mutations. A) A schematic shows various multicellular group shapes. These consist of an outer cell layer (red cells) and a pool of inner cells (gray cells). The fraction of inner to outer cells, *R*, determines the group’s ability to protect cells from stress, where higher values of *R* indicate more protection. For example we illustrate tightly packed spheres with *R* = 2.0, spheres with a half filled surface layer with *R* = 5.0, rod-shaped groups with *R* = 1.79, and biofilms with *R* = 0.78. B) A plot shows how beneficial mutations in the birth rate *b* have a larger positive impact on the fitness of populations of *U* (black solid line) compared to *M* (red dashed line). In the figure, relative fitness is compared to an ancestor with *b* = 1. C) A plot of the boundary from Eq. 2 that divides the parameter space into two regions, where either *U* (white) or *M* (red) has higher fitness. Beneficial mutations in *τ*_*G*_ and *b* enable populations to cross the boundary between the fitness regions. The type and order of mutations influence whether a population will maintain multicellularity (trajectory 2) or revert to unicellularity (trajectory 1). D) A plot shows the same boundary as in C), but for a higher *R* value. Increasing *R* provides an advantage to multicellular groups and expands the region where *M* is favored and can be maintained.

Similar to the case of differentiation, we can derive a criterion for when multicellularity (*M* ) has a fitness advantage over unicellularity (*U* ). The fitness advantage of *M* depends on how much protection is afforded to inner cells, which in turn depends on the length of time spent in the stressful (*E*_*A*_) environment. To simplify the analysis, we assume that the stress persists for enough time that the groups no longer contain inner cells (see Supplementary Information “Derivation of undifferentiatied multicellularity” for analyses of different stress durations). Based on this assumption, *M* has higher fitness than *U* when

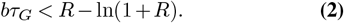

Here, *bτ*_*G*_ expresses the cost of being multicellular in terms of the missed growth potential when groups break apart. The cost of being multicellular must be offset by the protection afforded, which is represented by the *R*− ln(1 + *R*) term on the right hand side of Eq. (2). By reducing the time lag, populations can lower the cost of being multicellular; however, if populations increase the birth rate *b* then this favors unicellularity more than multicellularity, see Fig. 3B.

By constructing a parameter space for *b* and *τ*_*G*_ and calculating the relative fitness for *M* vs *U* we can again see how the order of beneficial mutations affects evolutionary trajectories. In the case where group protection is low, i.e. *R* is low, early mutations in the birth rate *b* can lead populations into the region where *U* is fittest. Since mutations in *τ*_*G*_ are neutral to *U* and may not fix, populations can get trapped in the unicellular life cycle, similar to the case with differentiation. If instead, early mutations occur in *τ*_*G*_ it is possible to lock populations into a region where multicellularity is fittest, see Fig. 3C. These trajectories are also shaped by the structure of the group, as higher values of *R* produce larger areas of parameter space (in terms of *b* and *τ*_*G*_) where *M* is favored, see Fig. 3D.

### Differentiated multicellularity

A final strategy for improving fitness during abiotic stress combines cell differentiation with multicellularity, to produce differentiated multicellularity (*M*_*D*_). While in principle there are many ways to combine differentiation with multicellularity (see Supplementary Information “Derivation of differentiated multicellularity” for other alternatives), we consider a way that minimizes interference and gives rise to complementary benefits. We assume that upon exposure to the abiotic stress cells first differentiate and then form groups. Since the cells on the exterior of the group are protected from the stress, they do not die and require replacement. This allows interior cells to de-differentiate so that when the stress disappears they can rapidly start growing once the multicellular group is disbanded — thus they do not experience the time lag for differentiation in the growth environment. When leaving the stressful environment, groups first dissociate. Afterwards, inner cells can start growing immediately, while outer cells first need to undergo differentiation. Thus, the differentiated multicellular lifestyle has the potential to harness the benefits of both differentiation and group formation, but it also incurs the combined costs. We can derive criteria for when differentiated multicellularity offers a fitness advantage compared to either differentiation alone (see Eq. (3)) or multicellularity alone (see Eq. (4)).

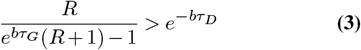

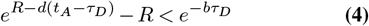

Since the resulting criteria contain more than two parameters, visualizations of the adaptive trajectories depend on the choice of parameters. In Supplmentary Information “Comparing fitness conditions for *U*_*D*_ vs *M*_*D*_” we consider all pairwise choices for parameters in Eq. (3) for *M*_*D*_ versus *U*_*D*_. These pairwise plots show that adaptation in some traits such as the growth rate *b* can have little effect on which strategy is fitter. Adaptation in other traits such as the differentiation time delay *τ*_*D*_ can cause all initial conditions favoring *M*_*D*_ to eventually enter a region where *U*_*D*_ is fitter.

In the case of differentiated multicellularity versus multicellularity the criterion determining which is fittest also depends on several parameters (see Supplementary Information “Comparing fitness conditions for *M* vs *M*_*D*_”). One difference with this criterion is that it draws a distinction between the growth rate *b* and the death rate *d*. We find that adaptation in *b* can lead to increased relative fitness of *M*_*D*_ while adaptation in *d* can have the opposite effect, eventually leading to an area of parameter space where *M* is fitter.

The conditions Eq. (3) and Eq. (4) for when differentiated multicellularity is fitter than its evolutionary antecedents share some trait parameters, such as the time lag for differentiation *τ*_*D*_. If we consider beneficial mutations that decrease *τ*_*D*_ we find different effects on the relative fitness of differentiated multicellularity compared to differentiation *U*_*D*_ and multicellularity *M* alone. Fig. 4A shows that decreasing the delay *τ*_*D*_ causes *U*_*D*_ to be fitter than *M*_*D*_, favoring a loss of multicellularity. In contrast, Fig. 4B shows that when *d* is not small (i.e. *d >* 0.35) decreasing the delay *τ*_*D*_ causes *M*_*D*_ to be fitter than *M*, favoring differentiation. These two results occur because decreasing the time delay for differentiation favors differentiation over multicellularity which in turn alters the relative fitness of *M*_*D*_ compared to *U*_*D*_ and *M* .

**Fig. 4.**
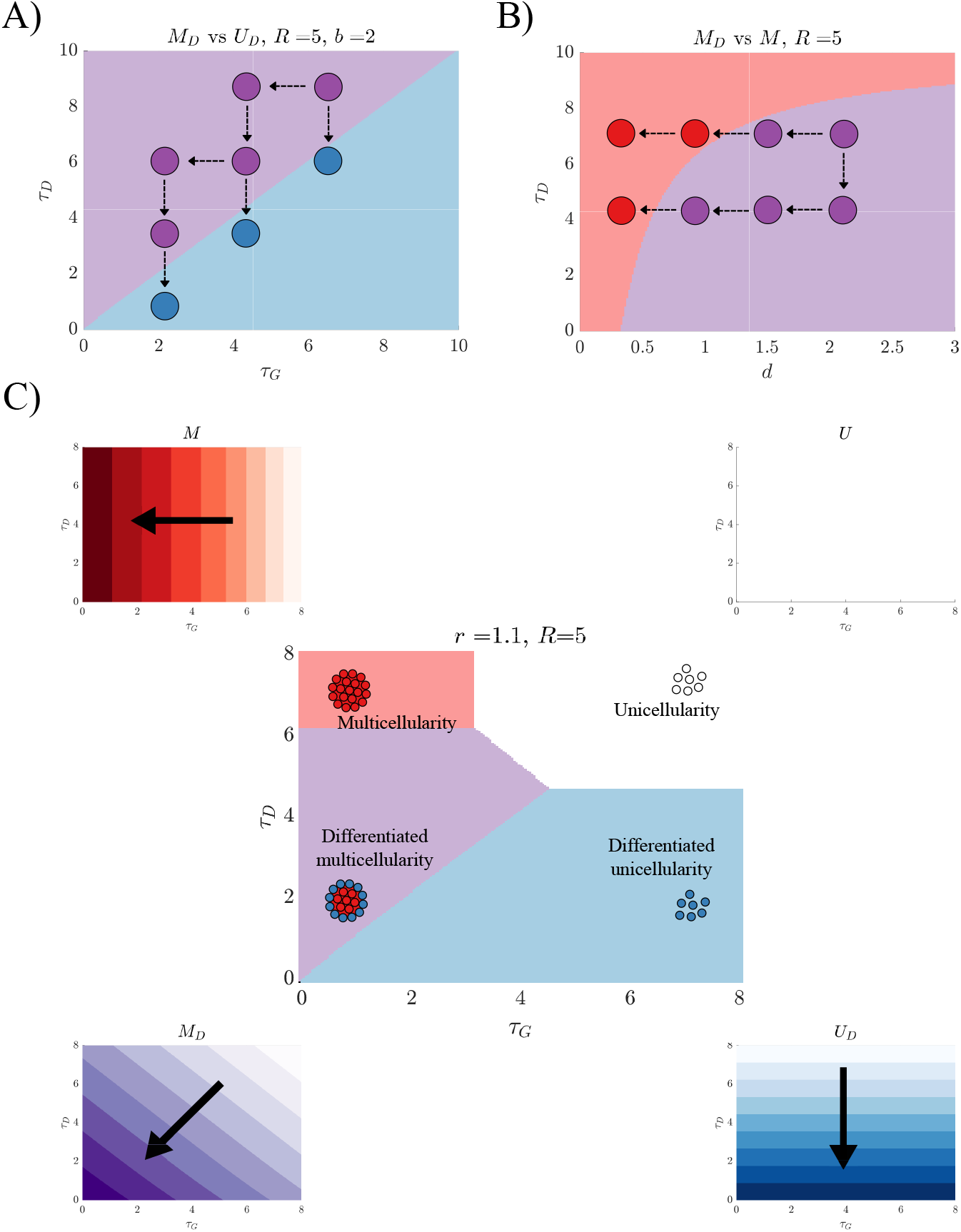
The many adaptive paths to and from differentiated multicellularity. A) A plot of the boundary from Eq. 3 in the *τ*_*G*_-*τ*_*D*_ plane. While beneficial mutations that decrease *τ*_*G*_ lead to entrenchment of the *M*_*D*_ phenotype (purple), mutations that decrease *τ*_*D*_ eventually lead to reversion to *U*_*D*_ (blue). B) A plot of the boundary from Eq. 4 in the *d*-*τ*_*D*_ plane. Beneficial mutations that reduce *d* eventually cause populations expressing *M*_*D*_ (purple) to revert to *M* (red), losing differentiation. In contrast to beneficial mutations that decrease *τ*_*D*_ reinforce *M*_*D*_ and prevent reversion to *M* . C) In the middle panel, a plot shows the *T*_*G*_-*T*_*D*_ (the time delays scaled by the time spent in the stressful environment) plane divided into regions where each phenotype has the highest fitness. The smaller surrounding panels illustrate how the fitness of each phenotype depends on *T*_*G*_ and *T*_*D*_ for example, *U* maintains the same fitness throughout the parameter space, while the fitness of *M*_*D*_ increases as either time lag is shortened.

We can combine the fitness calculations of the four phenotypes (*U, U*_*D*_, *M*, and *M*_*D*_) to identify which is fittest for a given combination of parameters. Fig. 4C shows the results of this calculation for parameters, *τ*_*G*_ and *τ*_*D*_ as well as the direction in parameter space where each phenotype increases fitness. Although each phenotype has a region where it is fittest, the regions for multicellularity and differentiated multicellularity are smaller than the region for differentiated unicellularity. If only beneficial mutations are considered then all phenotypes except for *M*_*D*_ are evolutionary stable in this parameter space. Differentiated multicellularity may remain the fittest provided the time delay for group formation is greater than differentiation, otherwise it will cross into a region where *U*_*D*_ is fittest.

### Evolutionary simulations

Until now we have considered adaptive evolutionary trajectories in regards to two traits at a time, but Eqs. 3-4 show that the relative fitness of differentiated multicellularity compared to multicellularity *M* or differentiation *U*_*D*_ depends on more parameters. To map possible adaptive trajectories in this higher dimensional parameter space, we perform evolutionary simulations that allow mutations in the defining traits of the life cycles (see Methods: Evolutionary simulations). More specifically, we consider mutations in the birth and death rates, the time delay for differentiation, and the time delay for group formation/dissociation. We also allow mutations to alter whether populations differentiate or form multicellular groups. Initially we only consider mutations that are beneficial and assume that they fix in a population with a probability based on the extent to which they increase relative fitness (see Methods: Evolutionary simulations). All simulations start with a unicellular life cycle and we simulate 10,000 independent evolutionary trajectories for different sets of initial parameters. Fig. 5 shows that for different sets of initial parameters we can bias evolution towards different types of complexity. For example by lowering the initial cost of differentiation we can increase the frequency of evolutionary trajectories that evolve differentiation, *U* to *U*_*D*_. Supplementary Information “Sensitivity to initial conditions” contains analyses for a more extensive set of initial parameters, which demonstrate that small changes in the initial values can alter the observed adaptive trajectories.

**Fig. 5.**
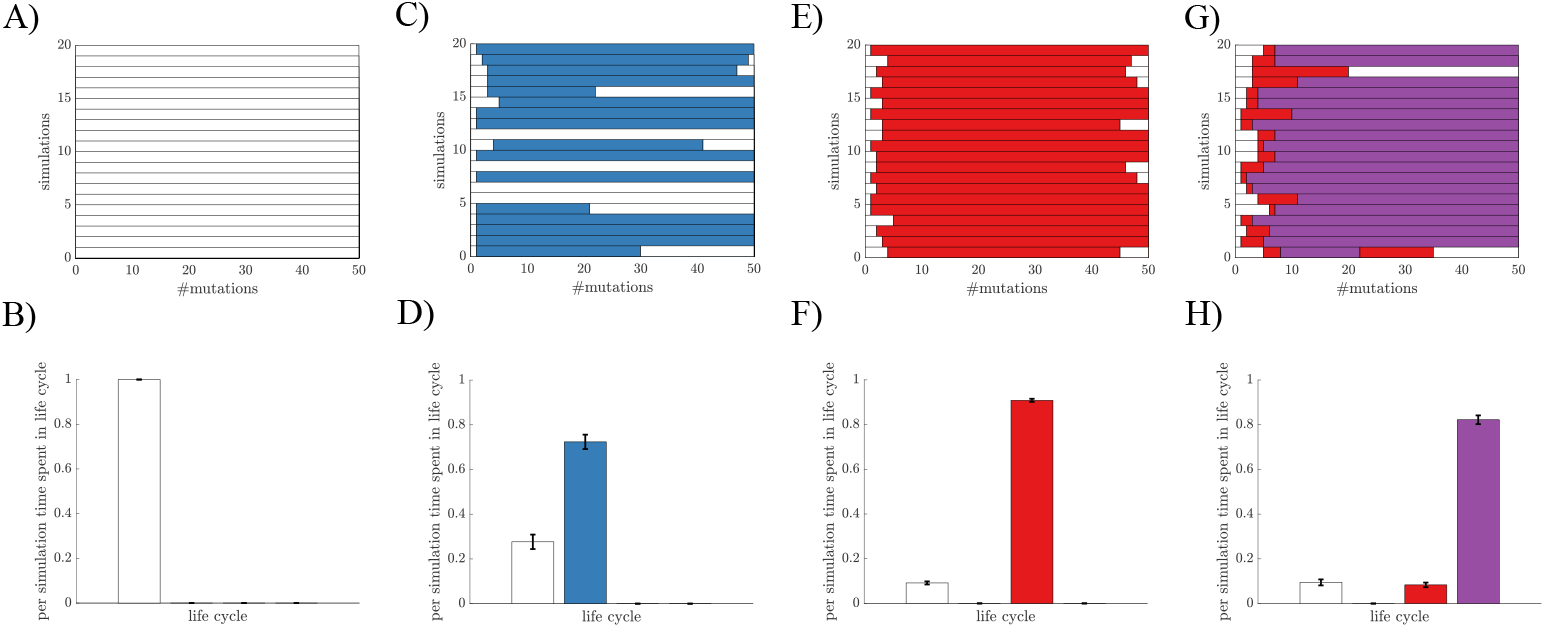
Historical contingency influences the evolution of complexity. A) A plot shows 20 sample evolutionary simulations. All mutations are adaptive and a mutation is accepted with a probability based on the relative fitness (see Eq. 9). The initial parameters for panels A-D) are listed in Supplementary Information “Sensitivity to initial conditions”. For the initial parameter setting used in A), only the *U* lifestyle (white) is observed. B) A histogram shows the fraction of time spent in each lifestyle; here, all time is spent as *U* . C) A plot shows the evolutionary outcomes for a different initial parameter setting, where some simulations result in the evolution of *U*_*D*_ (blue). D) A histogram shows that more time is spent in the *U*_*D*_ lifestyle than *U* . E)-F) Panels show similar data as C)-D), but for an initial condition that frequently results in the evolution of *M* (red). G) A plot shows the data from simulations with an initial setting where *M* (red) and *M*_*D*_ (purple) evolve. Since it takes two mutations, one in the ability to form groups and one to differentiate, we never observe a transition directly from *U* to *M*_*D*_. H) A histogram shows that *M*_*D*_ is most frequently observed given parameter setting in G).

The trajectories in Fig. 5 also indicate that there can be adaptive gains and losses of complex traits. By choosing different sets of initial parameters we can bias adaptation to evolve some form of complexity (*U*_*D*_, *M*, or *M*_*D*_) in over 99% of simulations (see Fig. 6A). Yet in 10-35% of the trajectories that evolved complexity, they ultimately reverted to unicellularity by the end of the simulation, see Fig. 6B. These losses of complexity occurred because although all mutations increased fitness, mutations in some traits, such as the death rate *d*, increased the fitness of unicellularity more.

**Fig. 6.**
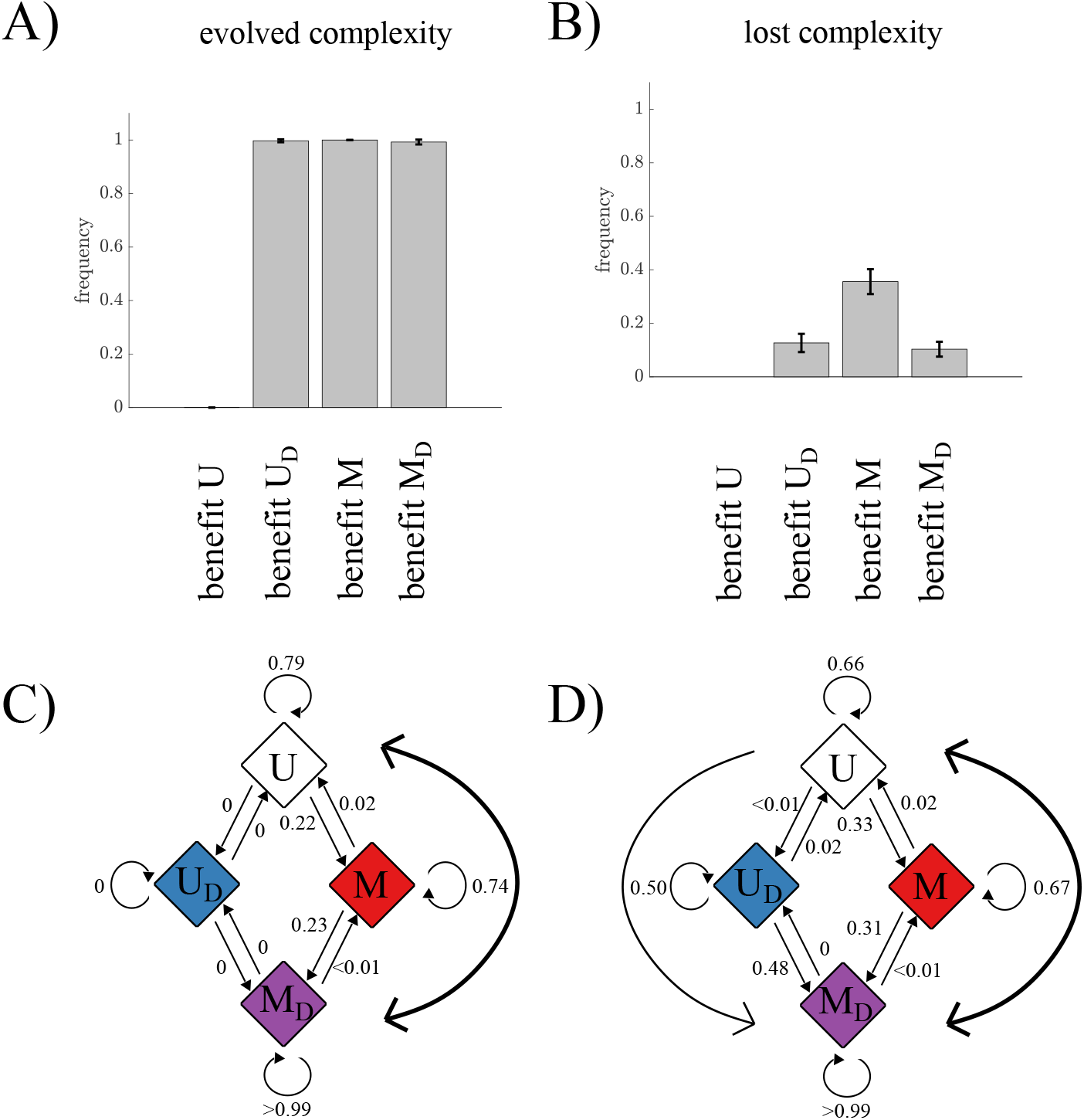
Gain and loss of complexity under constant selective pressure. A) A histogram shows the fraction of simulations where either differentiation or multicellularity evolved starting from each of the initial parameter settings used in Fig. 5. In over 99% of all simulations except those using the first set of parameters, some form of complexity evolved. B) A histogram shows the fraction of simulations that reverted back to *U*, after first evolving some form of complexity. Between 10 − 35% of simulations lost complexity and reverted back to unicellularity. C) A diagram represents the transition probabilities between phenotypes for the simulations in Fig. 5G. The data show that the most frequency path from *U* to *M*_*D*_ evolves multicellularity first. Similarly reversions from *M*_*D*_ to *U* happen by first losing the ability to differentiate. D) A diagram represents transition probabilities for the same case as in C), but neutral mutations are accepted with a 20% probability. Neutral mutations alter the evolutionary dynamics and here, the fraction of simulations that lead to evolution of *M*_*D*_ via evolving multicellularity *M* first.

We can use the data from the simulated evolutionary trajectories to determine the characteristic paths for gaining and losing complexity. Here we focus on the parameter setting that favored the evolution of differentiated multicellularity. In Fig. 6C we display the evolutionary paths between *U, U*_*D*_, *M*, and *M*_*D*_ as a state diagram where the arrows between states indicate transitions from one life cycle to another. Using this representation we see that when differentiated multicellularity evolves it typically proceeds from *U* to *M* to *M*_*D*_, i.e. it more often evolves multicellularity first rather than differentiation. Similarly, when differentiated multicellularity reverts back to unicellularity, it typically follows the same path in reverse, from *M*_*D*_ to *M* to *U* . We can alter the observed evolutionary paths from unicellularity to differentiated multicellularity by allowing neutral mutations to fix. Fig. 6D shows that when we permit neutral mutations to fix at a high rate, e.g. 20%, the trends from Fig. 6C are weakened as populations are freer to explore alternative paths to and from complexity (see Supplementary Information “Neutral mutations” for the affects of neutral evolution on other initial parameter settings).

These state diagram results pertain only to evolutionary simulations using a specific set of initial parameters. Supplementary Information section “Alternative evolutionary trajectories” shows state diagram representations for other initial parameter settings in which a majority of simulations evolved differentiated multicellularity. We find that for these settings the characteristic paths from *U* to *M*_*D*_ are different, going via *U*_*D*_ or both *U*_*D*_ and *M* . Thus historical contingency, represented here by the initial parameter settings, determines the characteristic paths towards complexity.

## Methods

### Equations for population growth

In order to assess the performance of each life cycle, we define fitness in terms of population growth. Fitness can be derived analytically under the assumption of exponential growth and death, and since the environment switches periodically between *E*_0_ and *E*_*A*_, it is sufficient to do the calculations over one such period. Starting with unicellularity *U*, we get

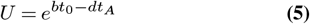

as the equation describing population fitness, where *t*_0_ is the time spent in *E*_0_ and *t*_*A*_ is the time spent in *E*_*A*_.

Fitness for differentiated unicellularity, *U*_*D*_, is calculated in a similar way as for *U*, except for a difference in the response to stress. We assume complete differentiation, meaning that when cells are in the survival state, there is no death in *E*_*A*_. However, since cells in this survival state cannot reproduce, there is a time lag *τ*_*D*_ when the environment switches back to *E*_0_ and cells must return to the growth state. During this lag cells do not reproduce, and so they miss an opportunity for population growth. Taken together the fitness is then

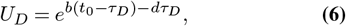

where the term 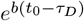 represents the growth in *E*_0_ and the term 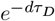 is the death that happens in *E*_*A*_ before the cells have differentiated.

For the multicellular life cycle, *M*, we need to consider the dynamics during the group phase in *E*_*A*_. As the outer cells of groups die and get replaced, the inner pool of cells will eventually be emptied. This happens at some time *t*^*^ in *E*_*A*_, and after this, the surface structure starts to collapse. Whether the time spent as a group in *E*_*A*_ has passed *t*^*^ affects the fitness, so we get two different expressions. We formulate them as

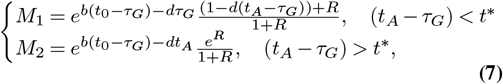

where the parameter *R* is the initial ratio of inner to outer cells in a group. The value of *R* serves as a measure of how well groups protect cells (see Supplementary Information “Derivation of undifferentiated multicellularity” for a full derivation). The time delay *τ*_*G*_ represents the mismatch of life cycle stage to environment, which happens when cells switch between single-celled and group state.

For the *M*_*D*_ life cycle, we assume that differentiation precedes group formation in *E*_*A*_, and multicellularity is lost before switching back to growth in *E*_0_. We note that other possibilities for the order of events exist but many involve interference between differentiation and multicellularity (see Supplementary Information “Derivation of differentiated multicellularity” for a review of the possible series of events). If we assume that groups are completely protected from stress and inner cells have de-differentiated so they can start to grow earlier in *E*_0_, we can formulate an expression for fitness:

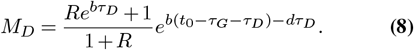

Similar to *M, R* is the ratio of inner to outer cells in groups. The fitness equations derived here can be used to make comparisons between the life cycles, see Supplementary Information “Pairwise comparisons”.

### Evolutionary simulations

We study the evolution of complexity in populations by simulating adaptive trajectories in which populations randomly fix beneficial mutations. Each simulation follows an iterative process in which mutations are chosen randomly from a uniform distribution. The mutations can alter the birth or death rates, *b* or *d*; the fraction of time used to switch between types, *T*_*D*_ = *τ*_*D*_*/t*_0_ or *T*_*G*_ = *τ*_*G*_*/t*_0_; or whether cells can differentiate or form multicellular groups. Initially we only consider beneficial mutations though we later expand to consider neutral mutations as well. Mutations that affect rates, e.g. *b* or *T*_*G*_, change the value by 5%. Each mutation fixes with a probability based on the amount by which it increases relative fitness, denoted as *f*_*rel*_, between the mutant and the resident type. We model the fixation probability to scale linearly with *f*_*rel*_ using the function:

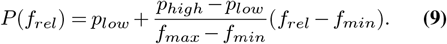

This implies that we accept all mutations with *f*_*rel*_ *> f*_*max*_ with the probability *p*_*high*_ and discard all mutations with *f*_*rel*_ *< f*_*min*_. Mutations within *f*_*min*_ *< f*_*rel*_ *< f*_*max*_ are accepted with a probability *p* in the range [*p*_*low*_, *p*_*high*_]. Specifically, the values used in the simulations are *f*_*min*_ = 1, *f*_*max*_ = 1.5, *p*_*low*_ = 0 and *p*_*high*_ = 1. This means that mutations resulting in a fitness benefit of more than 50% are always accepted, and all mutations must be beneficial to have a chance of fixing. For simplicity, we assume that beneficial mutations are rare, and so new mutations always appear in homogeneous populations.

Each simulation starts with as a unicellular *U* population. We follow the evolution until 50 mutations have fixed and repeat the process for 10,000 independent runs. The initial values of the time delays are shown in Fig. S5 and given in Tab. S1. Initial values for the other parameters are *b* = 1.1, *d* = 1, *t*_0_ = *t*_*A*_ = 10 and *R* = 5.

To study the effects of neutral mutations on adaptation, we adjust the parameters for the acceptance function in Equation 9. A neutral mutation has no effect on relative fitness, *f*_*rel*_ = 1, so to allow it a 20% chance of fixing we set *p*_*low*_ = 0.2. We note that mutations are neutral in regards to the current life cycle, so a mutation that is neutral may be deleterious should the life cycle change. For instance, a mutation in a unicellular population *U* that increases either of the time delays will be neutral but it would be deleterious in any of the other life cycles: *U*_*D*_, *M*, or *M*_*D*_.

## Discussion

Multicellularity has evolved independently in different lineages (3, 5, 14, 65, 66), and in many cases it has come with, or led to, further gains in complexity such as specialized cell types (13, 58, 64, 67, 68) or advanced body architectures (69– 71). Although there have been some investigations into the origins of complex multicellularity, they typically consider features in isolation, e.g. the evolution of different cell types within a multicellular organism. As a consequence, it is unknown whether the same pressure can lead to the evolution of complex forms of multicellularity from unicellular ancestors and what paths it might take. Here, we consider an abiotic stress as the main driver of selection and study the evolutionary paths between unicellularity and differentiated multicellularity. We use mathematical modeling and simulations to show that differentiated multicellularity can evolve from a unicellular ancestor in response to a single abiotic stress; however, the path to differentiated multicellularity is not permanent or guaranteed. Even when populations are exposed to a constant selective pressure and all mutations are beneficial, complex traits can be gained and lost adaptively, as populations continually improve their fitness. Thus, we find that differentiated multicellularity can evolve as a response to abiotic stress, but stress alone is not enough to stabilize a transition towards increased complexity.

One of the major findings of our study concerns the adaptive gains and losses of complexity. In our model cells experience regular bouts of abiotic stress. We observed gains in complexity because cells can improve their survival to the stress by either differentiating into stress-tolerant phenotypes or forming multicellular groups that offer protection. We observed losses of complexity because populations also experienced periodic bouts of no stress, where the only selective driver is growing as fast as possible. While all phenotypes benefit from mutations that increase growth rate, the unicellular phenotypes benefit the most because they do not spend time transitioning between forms, i.e. switching between unicellularity and differentiation/multicellularity. Once the growth rate is sufficiently high, the benefit of increased survival is not worth the cost of lost growth and so populations revert to unicellular forms. Consequently, the relative benefit of complex phenotypes versus unicellularity changes with the genetic background of the population, which means that populations can repeatedly gain and lose complexity in otherwise constant environments.

We expect these adaptive losses in complexity to be widely applicable to populations experiencing abiotic stress. For example, if the abiotic stress is an antibiotic then unicellular populations may be able to survive this stress by forming multicellular biofilms or differentiating into persistence states. If the population later evolves resistance to the stress— or alternatively the antibiotic is not particularly effective—then the relative benefits of evolving differentiation or multicellularity may not be worth their costs, driving populations to be unicellular. Whether such reversions occur will likely depend on a variety of factors including the costs of complexity, the genetic/phenotypic background of cells, and the limits to which populations can adapt in response to the stress. If, for example, the lag times impose a harsh limitation then the accumulated benefits of increasing growth rate (72) may lead to unicellular reversions. Since there are fundamental biological limitations in the initiation and elongation rates of transcription and translation it is not possible to switch phenotypes via changed gene expression and protein production faster than on the scale of minutes, which can be long enough for an antibiotic to kill most of the population (73). However, if populations can evolve ways of reducing the costs of complexity, e.g. the lag time in switching between phenotypes, then we expect complexity may be maintained. Unicellular organisms often use small molecule second messengers, including different nucleotides, Ca2+ and NO, to quickly respond to stress without needing to alter gene expression (74–78). Interestingly similar second messengers are also used as signals to form multicellular groups (79), including biofilm formation, as well as to initiate sporulation (74, 80); and they may have even been co-opted for the development of multicellular complexity (79, 81, 82).

Though we found that a single selective pressure could drive the evolution of differentiated multicellularity, whether it did so or not depended on two key factors: the initial phenotypic background and the order of mutations. The initial phenotypic background set the costs for forming multicellular groups and differentiating into protected types as well as the death/growth rates in the presence/absence of the abiotic stress. Thus, the initial phenotypic background determines how far a population is from a boundary where it is advantageous to change states, i.e. gain/lose differentiation and gain/lose multicellularity. From a given starting point, mutations that lowered the costs of multicellularity or differentiation acted to maintain forms of complexity or lead to further increases (i.e. evolving differentiated multicellularity), and mutations that increased growth rate or survival acted to promote reversions to unicellularity. The order of mutations also affected the evolutionary trajectories between unicellularity and differentiated multicellularity. In particular, we found that early mutations could lock populations into expressing complex traits, i.e. differentiation or multicellularity, and reduce the likelihood of reversions (see Fig. 2-4). If neutral mutations could fix then we found that differentiation and multicellularity more frequently evolved and reversions were less likely. This occurred because neutral mutations allowed unicellular populations to modify traits that increased the relative fitness of differentiation/multicellularity.

By studying the evolution of differentiation and multicellularity together within the same framework we were able to uncover ways in which these two types of complexity interfere with one another, inhibiting the evolution of differentiated multicellularity. One such example was in regards to the evolution of differentiation. Mutations that lowered the time to switch between stress-resistant and growth phenotypes favored the evolution of differentiated multicellularity *M*_*D*_ versus multicellularity *M* . In a framework where *M* and *M*_*D*_ are the only possibilities this would indicate that lowering the time to switch between phenotypes would lead to differentiated multicellularity. Yet, in another pairwise framework where *U*_*D*_ is compared with *M*_*D*_, the same type of mutation would lead to different conclusions. Lowering the cost to differentiation reduces the relative benefit of forming groups and favors unicellularity *U*_*D*_ over multicellularity *M*_*D*_. These results highlight that relying only on pairwise comparisons can constrain the evolutionary possibilities and miss important ways in which types of complexity interact.

In this study we considered a type of multicellularity in which groups form via aggregation, but there is another type of multicellularity in which cells remain in groups called clonal multicellularity. Both experimental and theoretical studies comparing these two types have found that clonal multicellularity may be better at gaining additional adaptations including more complex traits. In line with these findings, the majority of examples of large and complex multicellularity develop clonally (5, 14, 64, 83), yet there are some examples of aggregative multicellularity that have also evolved greater complexity such as cell differentiation and intricate multicellular structures (28, 84–86). For our study, we focused on aggregative multicellularity because it is typically found in systems that experience environmental fluctuations similar to those imposed by many abiotic stresses, although it has recently been discovered that some organisms can employ a combination of aggregation and clonality (16). By alternating between unicellular and multicellular forms, aggregative multicellularity can capitalize on the relative advantages in the different environments. In contrast, clonal multicellularity can offer survival benefits in response to stress but is at a disadvantage when the stress is removed and populations compete for rapid growth (63). Thus, we would expect clonal multicellularity to evolve in scenarios where the costs to cell reproduction are outweighed by the benefits accrued to surviving the abiotic stress. Since cells remain in groups in clonal multicellularity, there are more opportunities for the evolution of traits whose trade-offs limit reversion to unicellularity, e.g. those that lead to the creation of specialized cell types that do not actively contribute to group reproduction. We expect that distinctions between aggregative and clonal multicellularity may significantly affect the role of abiotic stress in shaping the path to further complexity and would be an interesting area for future study.

To constrain the parameter space of our model we made assumptions concerning the structure of the abiotic stress and the population’s response, but there are variations on these assumptions that could alter the results. For example, we assumed the abiotic stress occurred regularly and was detectable so that cells had sufficient time to benefit from implementing strategies such as differentiation or multicellularity. There are other types of abiotic stress, such as variation in access to nutrients, that are severe and unpredictable which might make it difficult for cells to respond via differentiated multicellularity (87–89). In these selective environments populations may evolve heterogeneous strategies such as bet hedging, e.g. in the social amoeba *Dictyostelium discoideum* and the bacteria *Myxococcus xanthus* (91) a fraction of the population remains as single cells (loners and peripheral rods) while others aggregate in response to an altered environment. Instead of switching between unicellularity and multicellularity, populations may also adopt mixed strategies in which multiple life cycles are maintained simultaneously in a population (57, 92, 93). Here mathematical models can be particularly useful in elucidating how the specific structure of the abiotic stress may shape the eco-evolution of unicellular/multicellular populations.

In this paper we considered abiotic stress whereby the selective driver does not change in response to the evolving populations. We found that while abiotic stress has the potential to select for more complex forms of multicellularity, there are often reversions arising from adaptations that mitigate the severity of the stress. Another possible type of selective driver for multicellularity may be biotic in origin which would then have the potential to co-evolve (49). Examples of such drivers include predation or parasitism which both have been found to lead to the evolution of types of complexity (48) including multicellularity (6, 9)—though we note that in the case of multicellularity the predators were specifically prevented from co-evolving with the prey. It is unclear whether the co-evolutionary process will have any effect on the evolution of complexity. For instance if a prey evolves to be multicellular in order to escape predation, the predator may adapt to specifically target the multicellular prey thereby selecting either for reversion of the prey back to unicellularity or alternatively increased multicellular size. Distinguishing between these possibilities can help determine if any drivers of multicellularity are more likely to result in the evolution of further complexity or if the process is inherently dominated by chance events.

## ACKNOWLEDGEMENTS

The work is supported by grant 2018-03630 from the Swedish research council (https://www.vr.se/english). EL received the reward. The funders did not play any role in study design, data collection and analysis, the decision to publish, or preparation of the manuscript. We also thank Lindi Wahl for fruitful discussions and commentary.

## Supporting information

### Derivation of undifferentiatied multicellularity

In the undifferentiated multicellular lifestyle, *M*, cells grow as single cells in *E*_0_ but can improve their fitness by aggregating into groups when the stress is present, *E*_*A*_. We assume that groups are tightly packed so that there is no space for cell reproduction in the group phase, and that groups are spherical and of equal size. Moreover we assume that it takes some time *τ*_*G*_ to form and dissociate groups, so when entering *E*_*A*_ the cells initially die with the rate *d* and when leaving *E*_*A*_ they are temporally prevented from growth, see Fig. S1.

In a group, cells can be divided into two classes: the outer cells that make up the surface of the group and the inner cells. Cells inside a group are physically protected from the stress, while cells on the group surface die with the rate *d*. When a cell on the surface dies it immediately gets replaced by a cell from the inside. This divides the population growth of *M* into two cases depending on whether groups have any inner cells left when leaving *E*_*A*_. Growth of the *M* lifestyle can be described as

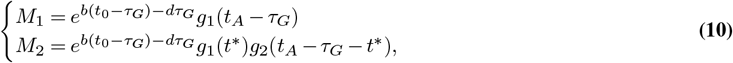

where *M*_1_ represents the case where groups still have inner cells left at the end of the stress phase, and *M*_2_ is the case when there are no inner cells left. The functions *g*_1_ and *g*_2_ describe the change in population size during *E*_*A*_ and *t*^*^ is the time it takes to empty the inner cell pool.

The functions *g*_1_ and *g*_2_ can be determined by modeling the dynamics of the inner and outer cells with a set of differential equations. Starting with the first case, *M*_1_, we can represent the dynamics for inner (*M*_1,*i*_(*t*)) and outer (*M*_1,*o*_(*t*)) as

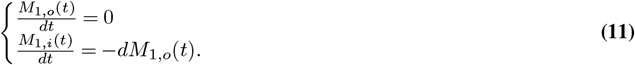

Integrating this gives

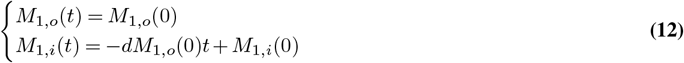

where *M*_1,*i*_(0) and *M*_1,*o*_(0) are the initial amounts of inner and outer cells. If *g*_1_(*t*) is the change in population size during *t*_*A*_ it can be expressed as

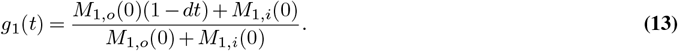

If we introduce 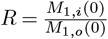 as the initial fraction of inner to outer cells and let *t* = *t*_*A*_ − *τ*_*G*_ be the time spent as groups in *E*_*A*_, we get the expression

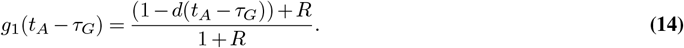

Putting this into *M*_1_ in Eq. 10 we get

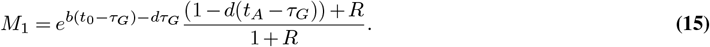

In order to derive *M*_2_ we first need to find an expression for *g*_2_, which represents the loss of outer cells when there are no inner cells left. We start by calculating how much time *t*^*^ it takes for the inner cell pool to be emptied. This is done by solving *M*_*i*,1_(*t*^*^) = 0 in Eq. 12 and we find that *t*^*^ = *R/d*. After this time the population dynamics changes in that the remaining outer cells just die with the rate *d* with no replacement of inner cells. So, when *t*_*A*_ *> τ*_*G*_ + *t*^*^ the population dynamics is described by

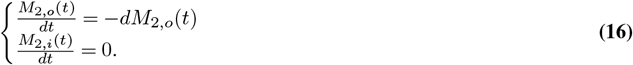

By integrating the upper equation we can express the change in population size as

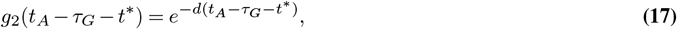

where *t*_*A*_ − *τ*_*G*_ − *t*^*^ is the time spent in *E*_*A*_ after the groups have run out of inner cells. Using that *t*^*^ = *R/d* we get

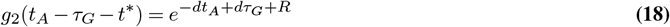

and putting this into the equation for *M*_2_ together with the result in Eq. 14 we get

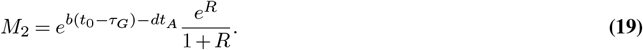

These calculations show that *M*_1_ and *M*_2_ can be expressed as

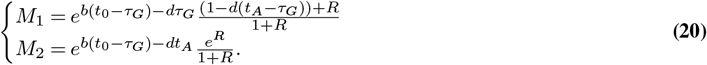

**Fig. S1.**
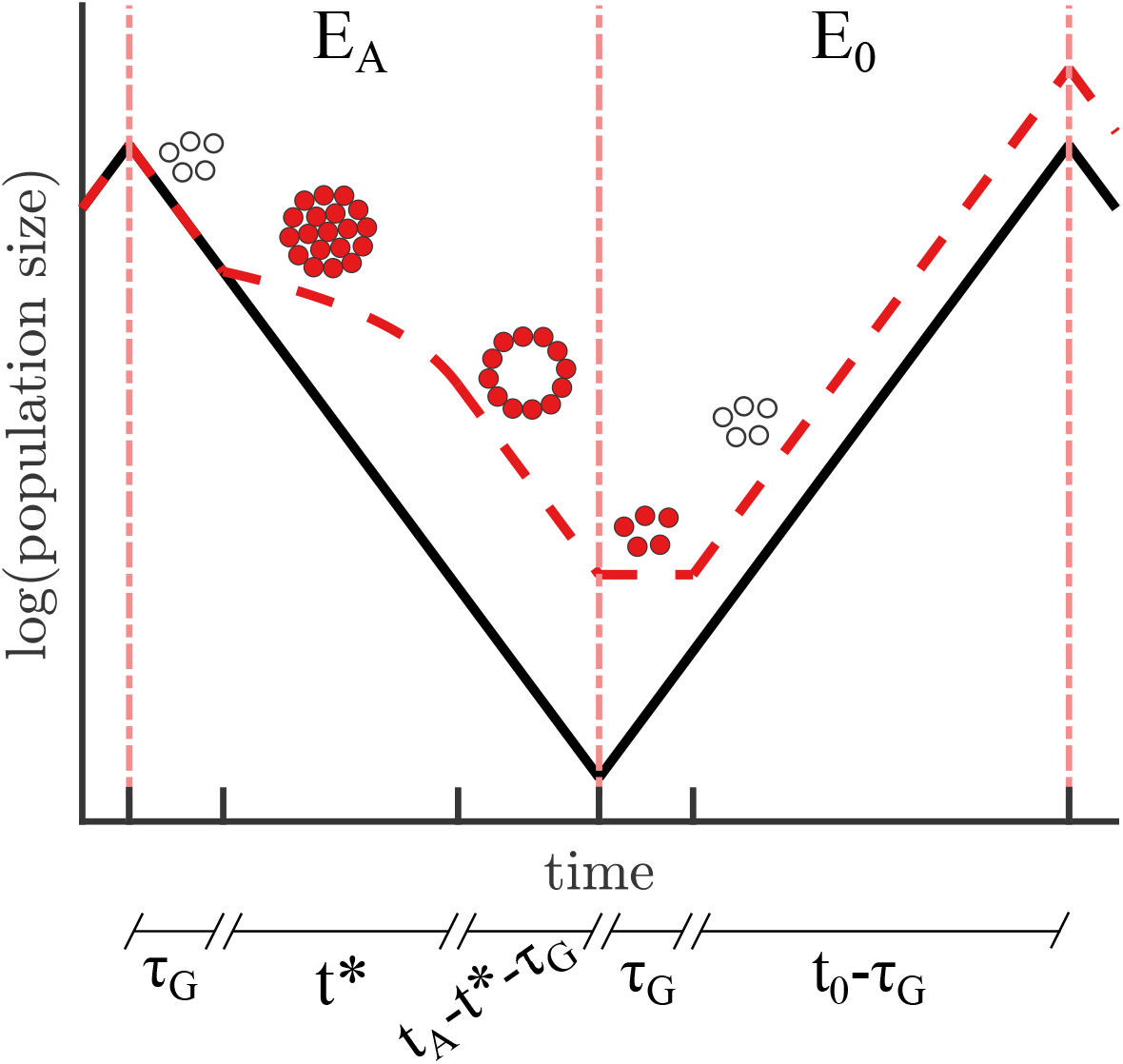
Multicellular groups provide physical protection during stress. A schematic illustrates the dynamics of two populations (*U*, solid line and *M*, dashed line) as the environment switches between *EA* and *E*_0_. The *U* population grows exponentially in *E*_0_ and dies in *E*_*A*_. The *M* population acts similar to *U* in *E*_0_, except that it initially experiences a time lag *τ*_*G*_ where it is prevented from growth. In *E*_*A*_, *M* dies at the same rate as *U* during *τ*_*G*_ when groups are formed. Once the groups are formed the death rate of *M* is reduced as long as there are inner cells left. When groups have run out of inner cells they die at the same rate as undifferentiated single cells.

### Derivation of differentiated multicellularity

Upon formation of differentiated multicellularity (*M*_*D*_) in *E*_*A*_ the outer cells are resistant to the stress and provide perfect protection for inner cells. Since inner cells are not affected by stress, they can switch back to a growth state while still in *E*_*A*_. This switching back to the growth state can provide an advantage in *E*_0_ allowing inner cells to grow as soon as multicellular groups have dissociated.

We can derive an expression for the fitness for *M*_*D*_, depending on the order of differentiation and group formation within the life cycle. Phenotypic switching can happen either sequentially e.g. groups can only form after all cells have switched to survival mode, or in parallel i.e. differentiation and group formation occur at the same time, see Fig. S2. The equations describing the fitness for each of the cases in Fig. S2 can be summarized as

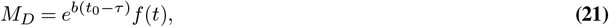

where the factor 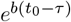 corresponds to population growth in *E*_0_ after groups have dissociated and cells have switched to growth mode. The function *f* (*t*) describes the change in population size during the phenotypic switching and it depends on the life cycle construction. There is no contribution from the population dynamics in *E*_*A*_ once cells have differentiated and formed groups since they would then have full protection from the stress.

If phenotypic switching happens in parallel we let *f* (*t*) = *f*_0,*D*_(*τ*_*D*_)*f*_0,*G*_(*τ*_*G*_)*f*_*A,D*_(*τ*_*D*_)*f*_*A,G*_(*τ*_*G*_) be a function that is composed of *f*_0,*D*_(*τ*_*D*_)*f*_0,*G*_(*τ*_*G*_) which corresponds to the population change during phenotypic switching in *E*_0_ and *f*_*A,D*_(*τ*_*D*_)*f*_*A,G*_(*τ*_*G*_) which corresponds to the population change during phenotypic switching in *E*_*A*_. For the case where switching between growth and survival modes always happens before group formation (*M*_*D*1_, first row in Fig. S2) we have 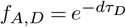 and *f*_*A,G*_ = 1 in *E*_*A*_ since all cells are protected after *τ*_*D*_. Similarly, in *E*_0_ we have that *f*_0,*D*_ = 1 and *f*_0,*G*_ = 1 because even if the inner cells have switched back to growth mode they need to wait until groups have dissociated. The resulting expression for *M*_*D*1_ is

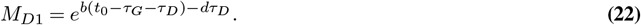

When group formation and dissociation precede switching between growth and survival modes (*M*_*D*2_, second row in Fig. S2) we let 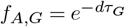 and 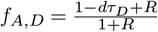 *E*_*A*_ (see Supplementary Information “Derivation of undifferentiated multicellularity” derivations for *M* ). Here, cells die with the rate *d* during group formation and while switching phenotype there is some death of the group’s outer cells. In *E*_0_, the inner cells can start to grow once the groups have dissociated while the outer cells first need to switch mode. Hence, *f*_0,*G*_ = 1 and 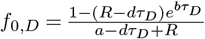 In particular, 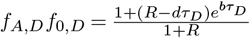 The final expression for *M*_*D*2_ is

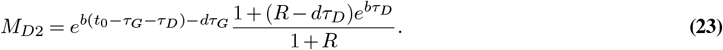

The *M*_*D*3_ case is when groups form before phenotypic switching in *E*_*A*_, and in *E*_0_ phenotypic switching happens first (third row in Fig. S2). This means that 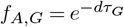 because cells die during group formation and 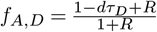 as some outer cells die before they have switched into survival mode. In *E*_0_ we have that *f*_0,*G*_ = 1 and *f*_0,*D*_ = 1 because there is no cell growth until the groups are dissociated. The resulting expression for *M*_*D*3_ is

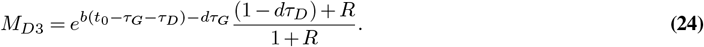

In the *M*_*D*4_ case (fourth row in Fig. S2) cells switch to survival mode in *E*_*A*_ before forming groups so 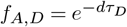 and *f*_*A,G*_*=1* as all cells have full protection during group formation. Upon removal of the stress (*E*_0_) the groups first dissociate so the former inner cells can start to reproduce while the outer cells also need to switch to growth mode. Particularly, this means that *f*_0,*G*_ = 1 and 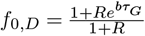 . To summarize, the growth equation for *M*_*D4*_ can be written as

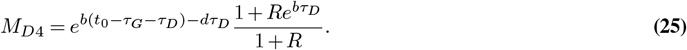

In both of the *M*_*D*5_ and *M*_*D*6_ cases the phenotypic changes happen in parallel. For *M*_*D*5_ (fifth row in Fig. S2) *τ*_*G*_ *> τ*_*D*_ so in *E*_*A*_ we have that 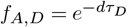 while *f*_*A,G*_ = 1. When the stress is removed cells are prevented from growth until the groups have dissociated so *f*_*A,D =*_1 while *f*_*A,G*_ = 1. In summary, the total growth of M_*D*5_ is

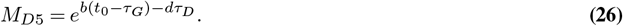

**Fig. S2.**
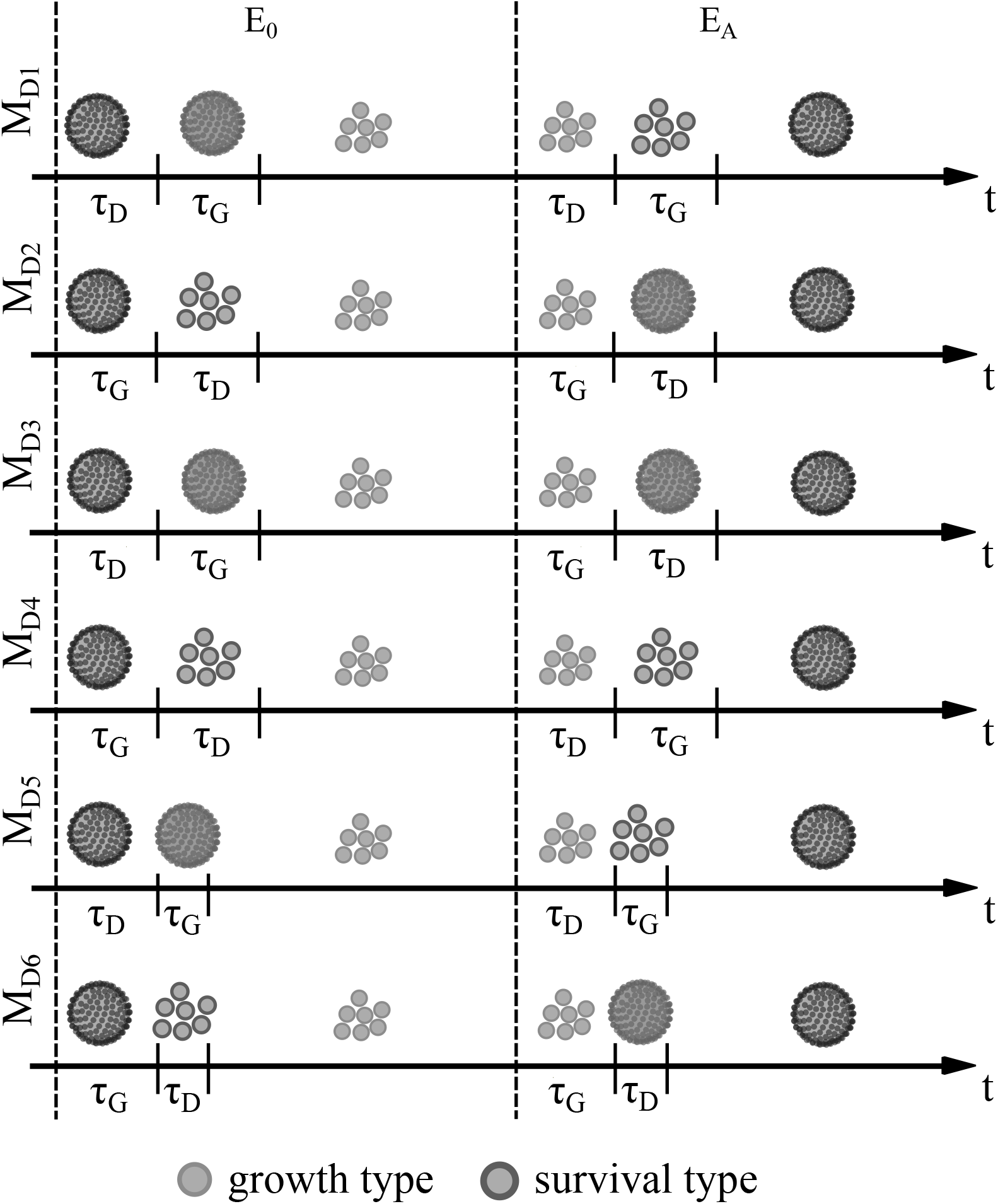
Effects from the order of phenotypic switches on the *M*_*D*_ life cycle structure. A schematic illustrates how the events, i.e. group formation/dissociation and switching between survival/growth phenotypes, can occur in different orders throughout the *M*_*D*_ life cycle. For all cases *M*_*D*_ enters *E*_*A*_ as undifferentiated single cells and leaves *E*_*A*_ as differentiated multicellular groups. In cases *M*_*D*1_ − *M*_*D*4_ the events are sequential i.e. one phenotypic switch happens at a time. For *M*_*D*5_ and *M*_*D*6_ the processes are parallel such that two phenotypic changes can take place simultaneously and the switching finishes faster. where Δ*τ* = *τ*_*D*_− *τ*_*G*_ in the expression for *M*_*D*6_. The *M*_*D*4_ case is the life cycle structure that we use for *M*_*D*_ in the main analysis.

The *M*_*D*6_ (sixth row in Fig. S2) is when *τ*_*D*_ *> τ*_*G*_. This means that in 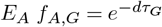 and 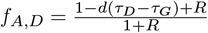 as there will be some death of the outer cells during the switch to survival mode. In *E*_0_ there is no cell reproduction during *τ*_*G*_ so *f*_0,*G*_ = 1, but after dissociation the inner cells can grow so 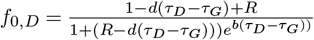. The total equation for fitness of *M*_*D*6_ is

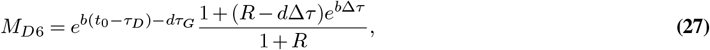

### Comparing fitness conditions for *U*_*D*_ vs *M*_*D*_

The fitness comparison between *U*_*D*_ and *M*_*D*_ depends on three variables (*τ*_*G*_, *τ*_*D*_ and *b*) so the there are several ways to construct a parameter space, see Fig. S3. From the figure we see that once populations are in a place where *U*_*D*_ is fittest it is not possible to evolve to a place where *M*_*D*_ is favored, assuming that only beneficial mutations are allowed. The reason is that the mutations required to evolve *M*_*D*_ are neutral to *U*_*D*_ so there is no selection for them to fix. In contrast, beneficial mutations in the *M*_*D*_ lifestyle, especially mutations in *τ*_*D*_, lead to loss of the multicellular trait. This analysis shows that by only allowing beneficial mutations it is difficult to maintain the *M*_*D*_ phenotype.

**Fig. S3.**
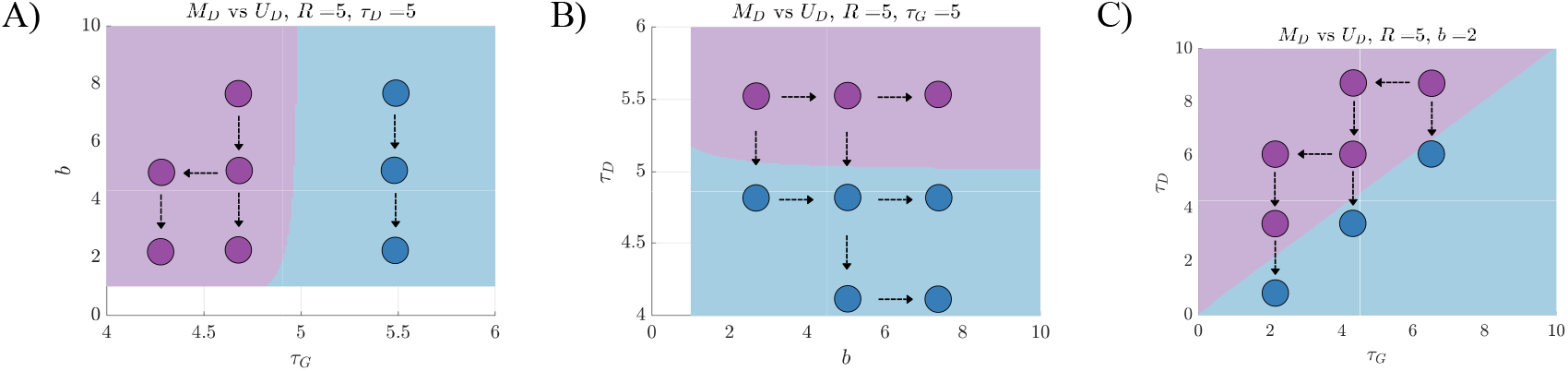
Beneficial mutations and reversions from *M*_*D*_ to *U*_*D*_. A) A parameter space shows which of *U*_*D*_ (blue) and *M*_*D*_ (purple) has the highest fitness with respect to the traits *τ*_*G*_ and *b*. Shown are two examples of adaptive walks in the parameter space where populations are locked into the *M*_*D*_ and *U*_*D*_ phenotypes respectively. B) A parameter space shows similar data as A), but for the traits *b* and *τ*_*D*_. Sample trajectories show that mutations in *b* mostly preserve the current phenotype while mutations in *τ*_*D*_ make *M*_*D*_ revert to *U*_*D*_. C) A similar parameter space as A) and B) but for the trait values of *τ*_*G*_ and *τ*_*D*_. Sample trajectories show that populations can maintain the *M*_*D*_ phenotype if mutations occur in *τ*_*G*_, but mutations in *τ*_*D*_ will lead to reversion to *U*_*D*_.

### Comparing fitness conditions for *M* vs *M*_*D*_

There are five parameters involved in comparing the fitness of *M* vs the fitness of *M*_*D*_: *R, t*_*A*_, *b, d*, and *τ*_*D*_. If we assume that *R* and *t*_*A*_ are not evolvable e.g. they are environmental variables, this leaves three parameters. The results from the comparisons are shown in Fig. S4, where the parameter space in panel A) shows that beneficial mutations in *b* and *τ*_*D*_ will change whether differentiation is fitter or not, except when *b* is very low. Adaptation in *b* affects *M* and *M*_*D*_ similarly so it does not change which is fitter. For the differentiation time delay, lower values of *τ*_*D*_ make *M*_*D*_ fitter, but mutations in *τ*_*D*_ are adaptive only in *M*_*D*_, because they are neutral in *M* . The parameter space in B) shows that improving survival to the stress, i.e. decreasing *d*, will eventually lead to loss of differentiation regardless of the value of *τ*_*D*_. Beneficial mutations in *τ*_*D*_ will lead to an area of parameter space where *M*_*D*_ is fitter as long as the death rate *d* is above a certain threshold *d* ≈ 0.32 here. Similar to the result in Fig. S3, the comparison of *M* and *M*_*D*_ shows that differentiated multicellularity is difficult to maintain when all mutations are beneficial.

**Fig. S4.**
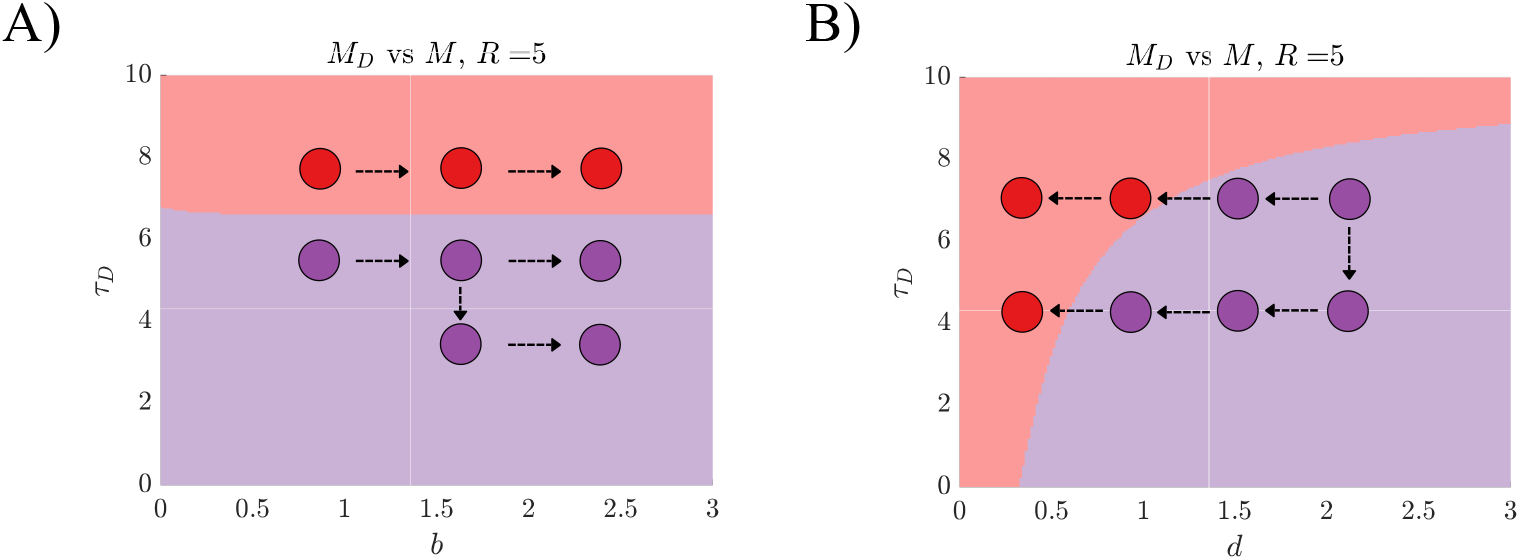
Beneficial mutations and reversions from *M*_*D*_ to *M* . A) A parameter space shows which phenotype *M* (red) or *M*_*D*_ (purple) is fitter for different combinations of traits *b* and *τ*_*D*_. Sample adaptive walks show that beneficial mutations in this parameter space rarely lead to transitions between the red *M* and purple *M*_*D*_ regions. B) A similar plot as in A) is shown but for the traits *d* and *τ*_*D*_. Here, the example adaptive walks show that trajectories stay in the *M*_*D*_ region for mutations in *τ*_*D*_, but mutations that reduce *d* will always lead trajectories into regions that only favor *M* .

### Sensitivity to initial conditions

For our simulations we selected initial conditions spread across parameter space that still allowed for adaptation (shown in Fig. S5 and listed in Table S1). We ran simulations for each set of initial conditions and tracked the phenotype (*U, U*_*D*_, *M*, or *M*_*D*_) in time. (see Fig. S6). We find that the initial condition largely dictates the evolutionary trajectory. In cases such as A) and J) the *U* phenotype occurs exclusively, because simulations that start in the *U* region cannot gain beneficial mutations in *τ*_*D*_ and *τ*_*G*_ which are needed to transition to regions that favor *M* or *U*_*D*_. The initial conditions used in Fig. 5 are A), C), F), and I) for panels A), C), E), and G) respectively.

**Fig. S5.**
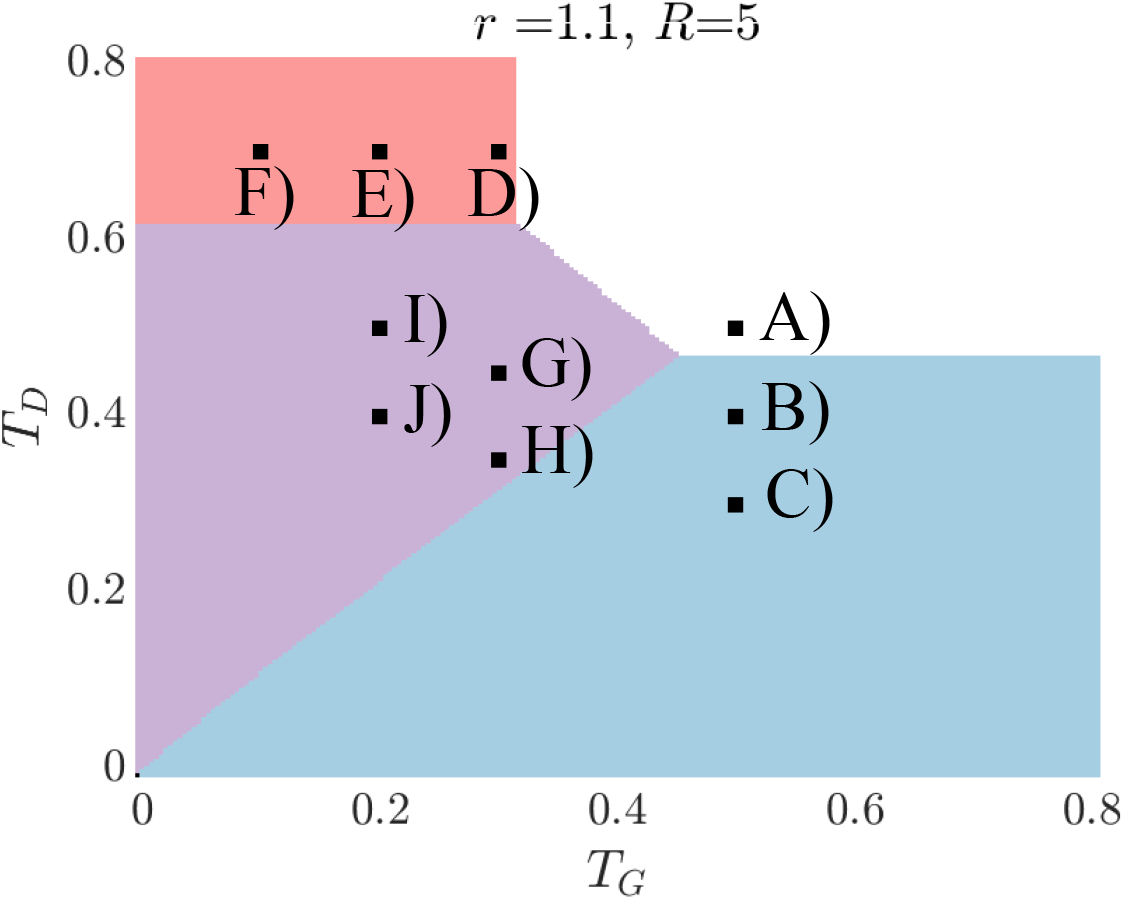
Initial parameter settings used for simulations. A set of initial conditions used for our simulations are shown in a parameter space. We use initial conditions from all four regions where each of the lifestyles has the highest fitness.

**Table S1.** Initial parameter values used for evolutionary simulations. Presented are the initial values of *τ*_*G*_ and *τ*_*D*_ (here normalized with the time spent in each environment) used for the evolutionary simulations.

| Initial position | $T_G$ | $T_D$ |
| --- | --- | --- |
| A) | 0.5 | 0.5 |
| B) | 0.5 | 0.4 |
| C) | 0.5 | 0.3 |
| D) | 0.3 | 0.7 |
| E) | 0.2 | 0.7 |
| F) | 0.1 | 0.7 |
| G) | 0.3 | 0.45 |
| H) | 0.3 | 0.35 |
| I) | 0.2 | 0.5 |
| J) | 0.2 | 0.4 |

**Fig. S6.**
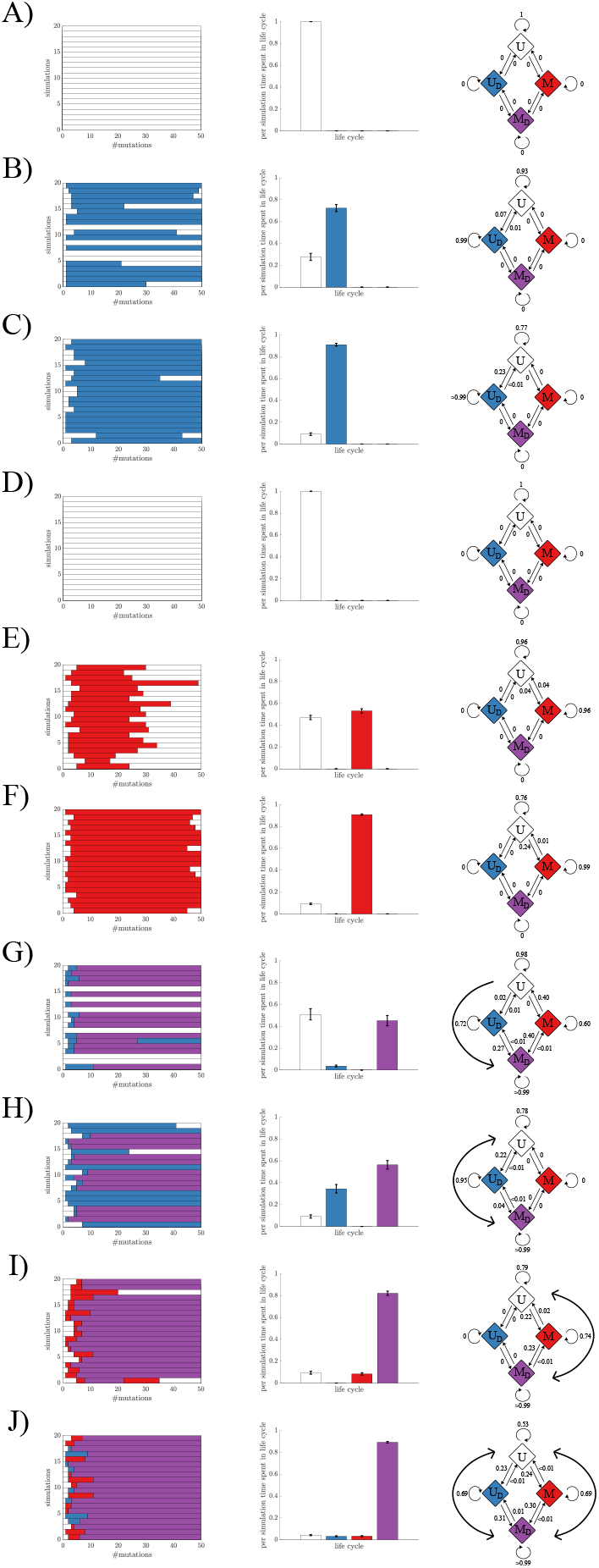
Initial parameter settings affects evolutionary transitions. A) A plot (left) shows a sample from simulations run for the initial condition A) in Fig. S5. The histogram (middle) shows the fraction of time during the simulations that is spent in each of the lifestyles. The diagram (right) shows the transition probabilities between the lifestyles. Panels B)-J) show similar data as panel A) but for the initial conditions B)-J) in Fig. S5.

### Neutral mutations

In the majority of our study we consider the fixation of beneficial mutations, i.e. those that increased fitness. Here, we allow mutations that are neutral to fix with a 20% probability. We note that neutral mutations do not change the fitness of the current life cycle but may have effects on fitness if the population evolves a different life cycle. An example of a neutral mutation is if a time delay in either differentiation or group formation changes in a *U* population. Allowing neutral mutations to fix opens up new evolutionary trajectories for each initial parameter setting, see Fig. S7. Neutral mutations also lead to higher frequency of gained complexity and less reversions, see Fig S8.

**Fig. S7.**
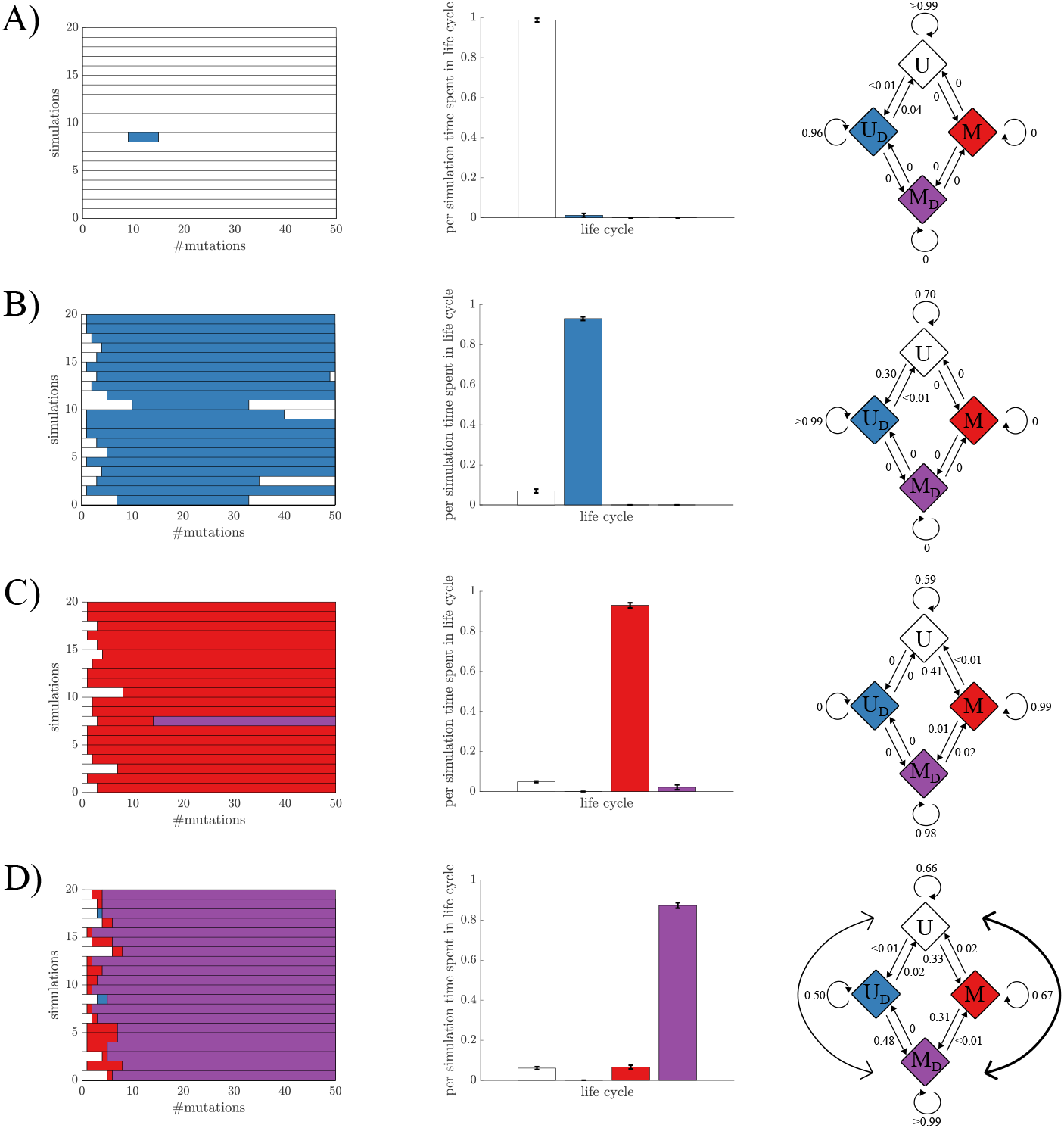
Neutral mutations and increased diversity of evolutionary routes. Neutral mutations are allowed with 20% probability, which opens up for evolution of new phenotypes. A) A plot (left) shows a sample from simulations run for the initial condition A) in Fig. S5 and here *U*_*D*_ is able to evolve with low frequencies. The histogram (middle) shows the fraction of time during the simulations that is spent in each of the lifestyles. The diagram (right) shows the transition probabilities between the lifestyles. B) The panels show similar data as A) but for the initial condition C) in Fig. S5. Since *U*_*D*_ is already dominant in this setting, neutral mutations do not have a big impact on evolution. C) The panels show similar data as A) but for the initial condition F) in Fig. S5. Allowing neutral mutations enables the *M*_*D*_ lifestyle to evolve in this setting. D) The panels show similar data as A) but for the initial condition I) in Fig. S5. The neutral mutations open up for new evolutionary routes from *U* to *M*_*D*_, in this case from *U* to *U*_*D*_ to *M*_*D*_.

**Fig. S8.**
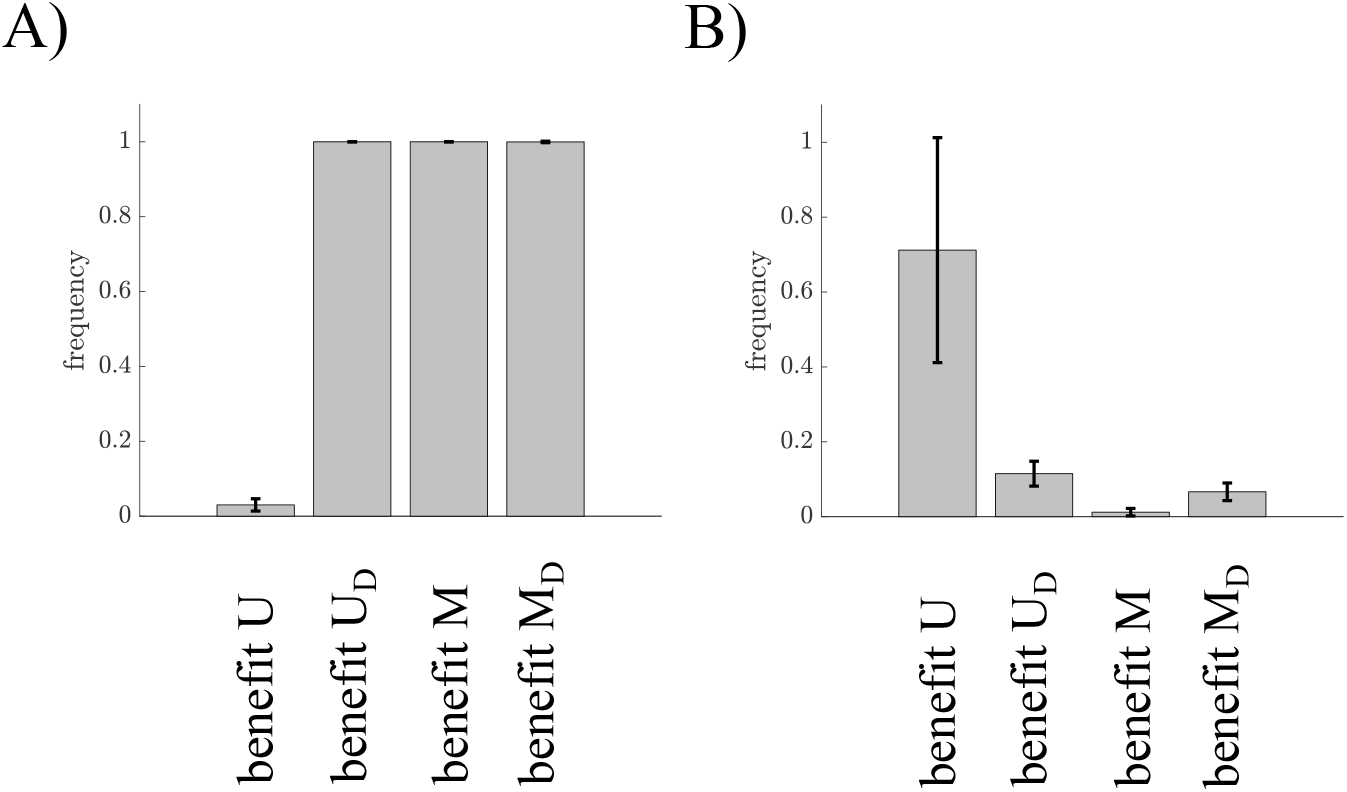
Neutral mutations enhance evolution of complex lifestyles. A) A histogram shows the frequency of evolved complex traits when neutral mutations are allowed. Compared to Fig. 6A) complex traits evolve to a larger extent when neutral mutations are allowed. A histogram shows the frequency of reversions back to *U* given that complexity first evolved. We observe that reversions are more rare than when mutations are only beneficial, see Fig. 6B). Here, *U* is an exception because for beneficial mutations this case did not evolve complex traits at all.

### Alternative evolutionary trajectories

The routes from *U* to *M*_*D*_ can be altered by varying the initial parameter settings. By starting the simulations with trait values close to the *U*_*D*_ boundary, such as case H) in Fig. S5, the evolutionary trajectory goes from *U* to *U*_*D*_ to *M*_*D*_. For a different starting point we find that evolution takes other paths. Using case J) in Fig. S5 as the initial state, *M*_*D*_ can be reached via either of *U*_*D*_ first or *M* first.

**Fig. S9.**
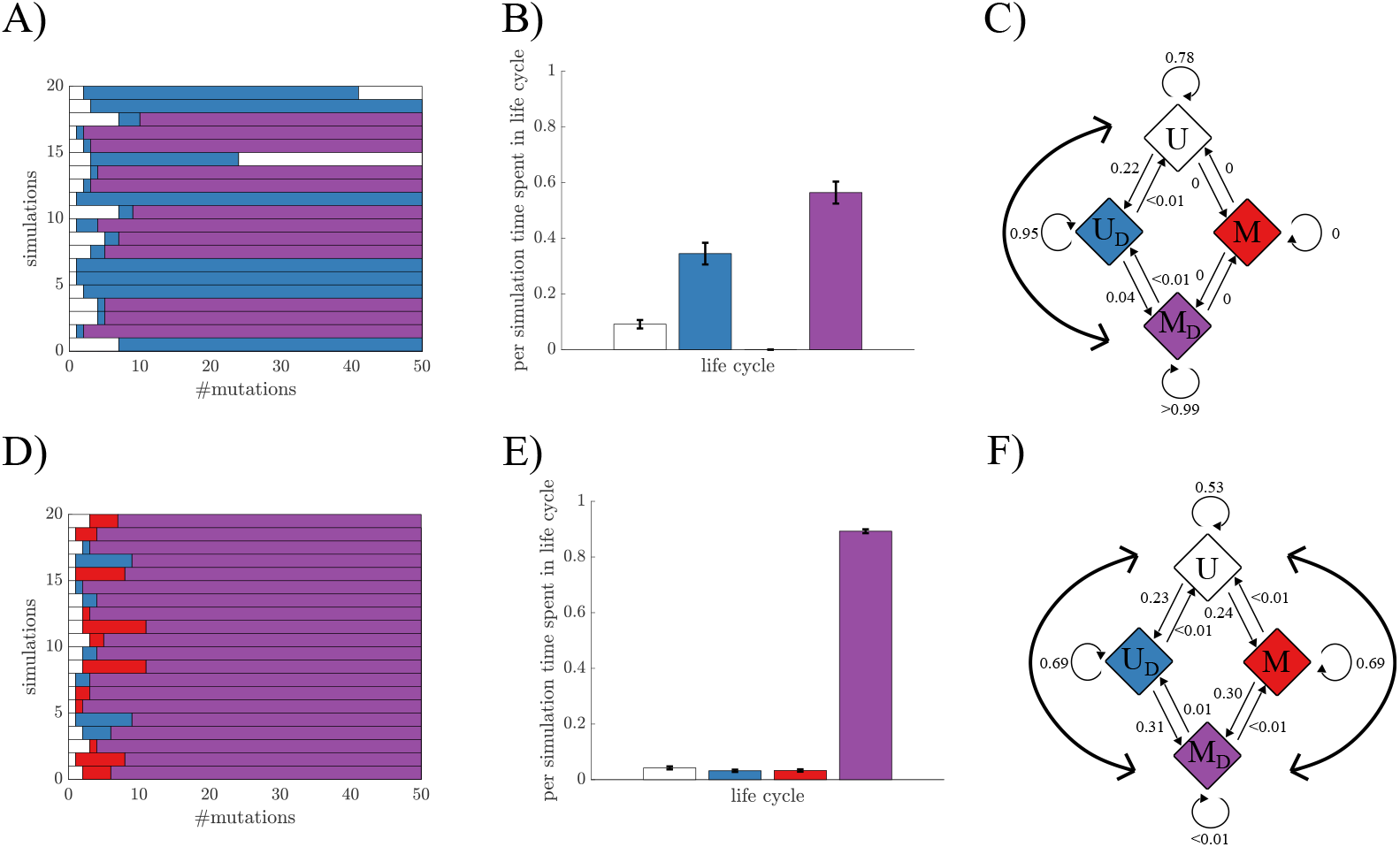
Evolutionary trajectories take different routes towards differentiated multicellularity. A) A plot shows a sample from simulations where H) in Fig. S5 was used as the initial condition. The data shows that *M*_*D*_ evolves via a transition to *U*_*D*_. B) A histogram shows the fraction of time spent in each lifestyle, and here *M*_*D*_ is the most common. C) A diagram shows the transition probabilities between the lifestyles. For the initial condition H) the route between *U* and *U*_*D*_ goes via *U*_*D*_. D)-F) Similar data is shown as in A)-C) but for the initial condition J). In this case evolution can take either of the routes i.e. differentiation first or multicellularity first, from *U* to *U*_*D*_.

### Pairwise comparisons

The relative fitness of life cycles can be analyzed by doing pairwise comparisons. Here we derive analytical expressions for the conditions when one life cycle is fitter than another.

### Differentiated unicellularity vs undifferentiated unicellularity

First we derive a condition where differentiated unicellularity *U*_*D*_ is fitter than undifferentiated unicellularity *U* . Taking the logarithms of the growth equations (Eq. 5 and Eq. 6) and setting *U*_*D*_ *> U* we get

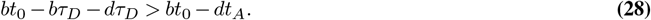

Dividing by *d* and *t*_*A*_ we can write the expression in a dimensionless form. If we define the dimensionless parameters as

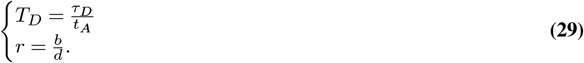

and solve for *T*_*D*_ the resulting condition for when *U*_*D*_ has higher fitness than *U* is

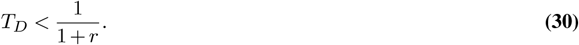

### Undifferentiated multicellularity vs undifferentiated unicellularity

With a similar approach we can calculate when undifferentiated multicellularity *M* is fitter than undifferentiated unicellularity *U* . For *M* there are two cases depending on whether any inner cells are left (see Supplementary Information “Derivation of undifferentiated multicellularity”). Here, we assume that the stress is around long enough for groups to run out of inner cells. Using Eq. 5 and Eq. 7 and taking the logarithm of *U < M* we get

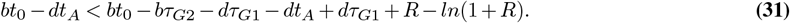

Simplifying this gives

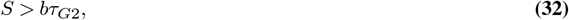

where *S* = *R* − ln (1 + *R*) is the shape parameter that depends on the fraction of inner to outer cells (*R* = *M*_*i*_(0)*/M*_*o*_(0)).

### Differentiated unicellularity vs differentiated multicellularity

For a comparison between *U*_*D*_ and differentiated multicellularity (*M*_*D*_) we use Eq. 6 and Eq. 8. Taking *U*_*D*_ *< M*_*D*_ and rearranging terms gives the condition

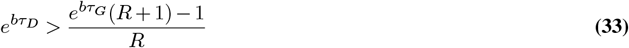

for when *M*_*D*_ has higher fitness.

### Undifferentiated multicellularity vs differentiated multicellularity

Comparing under what conditions *M*_*D*_ is fitter than *M* we use Eq. 8 and Eq. 7 and again make the assumption that *M* has run out of inner cells when leaving *E*_*A*_. By setting *M < M*_*D*_ and rearranging terms we get the expression

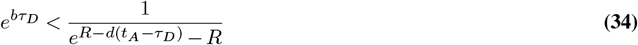

for when *M*_*D*_ has higher fitness than *M* .

